# A Cancer-Associated Fibroblast Classification Framework for Single-Cell Data

**DOI:** 10.1101/2022.12.14.520398

**Authors:** Lena Cords, Sandra Tietscher, Tobias Anzeneder, Claus Langwieder, Martin Rees, Natalie de Souza, Bernd Bodenmiller

**Affiliations:** Department of Quantitative Biomedicine, University of Zurich, CH-8057 Zurich, Switzerland; Institute of Molecular Health Sciences, ETH Zurich, CH-8093 Zurich, Switzerland; Life Science Zurich Graduate School, ETH Zurich and University of Zurich, CH-8057 Zurich, Switzerland; Patients’ Tumor Bank of Hope (PATH), D-81337 Munich, Germany; Pathology at Josefshaus, D-44137 Dortmund, Germany

## Abstract

Cancer-associated fibroblasts (CAFs) are a diverse cell population within the tumour microenvironment, where they have critical effects on tumour evolution and patient prognosis. To define CAF phenotypes, we analysed a single-cell RNA sequencing (scRNA-seq) dataset of over 16,000 stromal cells from tumours of 14 breast cancer patients, based on which we defined and functionally annotated nine CAF phenotypes and one class of pericytes. We validated this classification system in four additional cancer types and used highly multiplexed imaging mass cytometry on matched breast cancer samples to confirm our defined CAF phenotypes at the protein level and to analyse their spatial distribution within tumours. This general CAF classification scheme will allow comparison of CAF phenotypes across studies, facilitate analysis of their functional roles, and potentially guide development of new treatment strategies in the future.

## Introduction

The tumour microenvironment (TME) is a complex ecosystem consisting of diverse and interacting cell populations. These cell populations include a variety of resident and infiltrating immune cells and stromal cells as well as tumour cells. The composition of the TME influences tumour progression and metastasis^1^, the anti-tumour immune response^2^, and therapy response^3^. It is thus crucial to study tumours as complex ecosystems in order to understand intercellular interactions and to improve patient prognosis.

Fibroblasts are the main constituent of the tumour stroma. Cancer-associated fibroblasts (CAFs) are diverse cells with numerous roles within the TME^4^. They are key players in shaping the tumour microenvironment with functions in tumour promotion^5, 6^ and inflammation^7–9^ as well as maintenance and reshaping of the extracellular matrix (ECM)^4, 10^. Different CAF subpopulations have been described in various cancer types^11–14^. In breast cancer, for example, single-cell RNA sequencing (scRNA-seq) in a mouse model detected four different CAF phenotypes, termed vascular CAFs, matrix CAFs, cycling CAFs, and developmental CAFs^15^. A separate scRNA-seq study using a triple-negative breast cancer mouse model further identified CAF phenotypes with inflammatory and immune regulatory, ECM producing, protein folding, and antigen-presenting functions^16^. In human breast cancer, the CAF-S1 subtype is associated with an immunosuppressive environment^13^ and cancer cell migration, thus likely promoting metastasis^17^.

Despite recent advances, there is still no decisive set of markers for unambiguous identification of CAFs and CAF subtypes in the TME^10^. Although CAF subsets have been identified and linked to patient prognosis or therapy response^18^ in pancreatic ductal adenocarcinoma (PDAC)^7, 9^, bladder urothelial cancer^14^, melanoma^19^, and lung cancer^11, 18^, the use of small cohorts or limitations of available markers mean that the identified CAF subsets are typically studied only per cancer type^18, 20, 21^. A few studies have evaluated marker expression in CAFs across species^7, 9, 12, 22^ and across cancer types^22^, but it remains unclear whether defined CAF phenotypes are widely applicable. To gain a comprehensive understanding of CAF heterogeneity, a tumour type-independent CAF classification system is needed.

Here, we analysed a scRNA-seq dataset of human breast cancer and defined a total of ten distinct CAF and one pericyte populations (Figure 1). We confirmed these populations at the protein level using multiplex imaging mass cytometry (IMC) on matched samples, which also allowed us to investigate their spatial distribution within the TME and confirm our marker-based classification system (Figure 1). Our analysis across several tumour types showed that these cellular subpopulations are present across cancer types and thus that the defined CAF phenotypes can be generalized. In summary, we propose a general classification system for CAFs based on gene expression profiles and define their spatial features such as distance to the stroma/tumour border, to structures such as vessels, and to classes of neighbouring cells (Figure 1).

**Figure 1.**
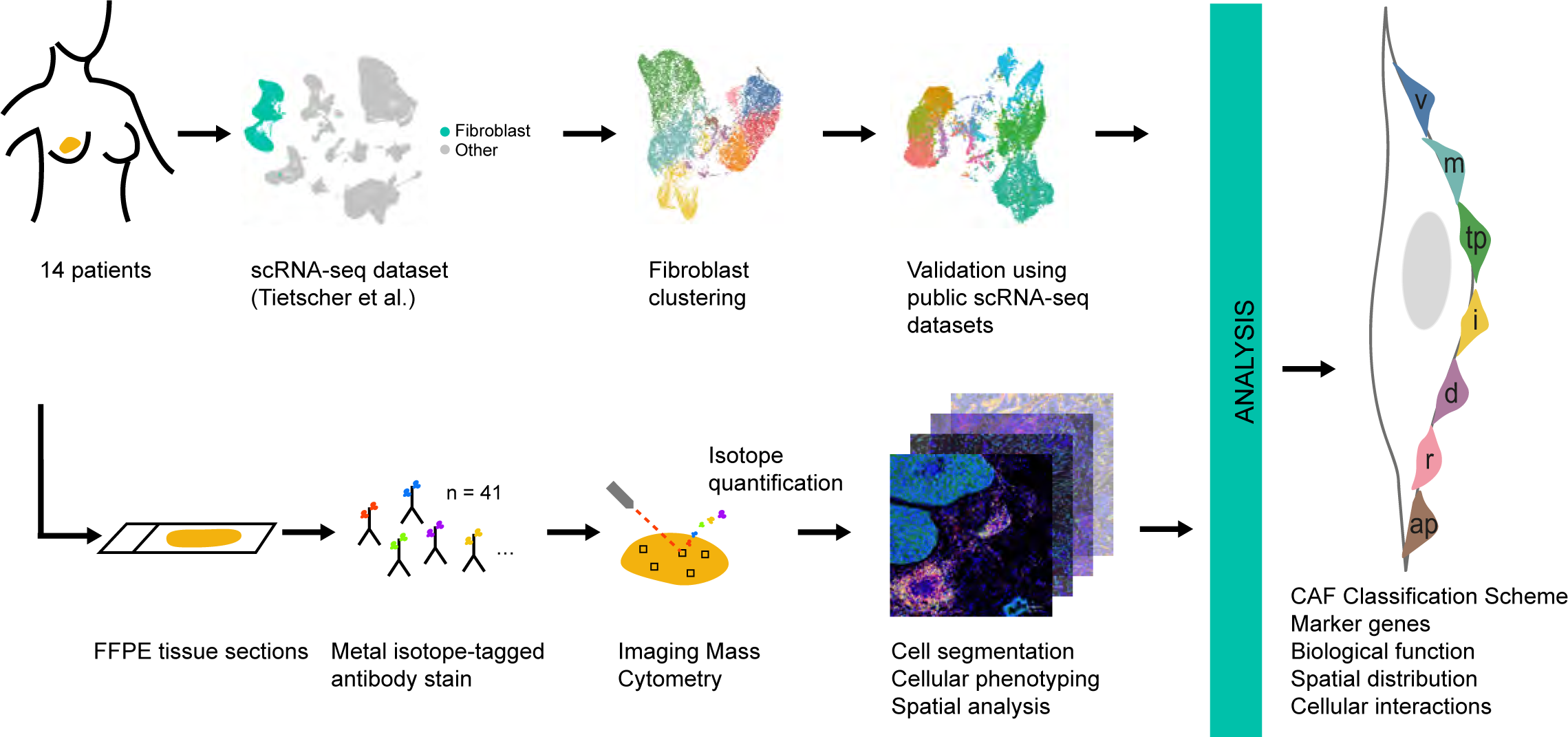
Workflow used to define the CAF classification system. scRNA-seq data from matched breast cancer samples (Tietscher et al., in press) were analysed for fibroblast heterogeneity with subsequent validation of identified fibroblast subtypes in other tumour types. IMC was used to validate findings at the protein level and to evaluate spatial distribution. The resulting CAF classification system was based on marker genes, biological functions, spatial distribution within the TME, and cellular interactions.

## Results

### Fibroblast classification in breast cancer

To identify and classify CAF types, we analysed our previously generated scRNA-seq dataset from 14 human breast cancer specimens (Supplementary Table 1), in which we had identified 16,704 stromal cells (out of ∼119,000 cells total) based on unsupervised clustering (Tietscher et al., in press). Of these stromal cells, we identified 2,389 cells as pericytes based on expression of *RGS5*^23^, resulting in a total of 14,315 CAFs. We used this data set to investigate fibroblast phenotypic heterogeneity in breast tumours.

After batch correction (Supplementary Figure 1a), we performed unsupervised hierarchical clustering of the single-cell gene expression profiles of all stromal cells, which identified 12 clusters at a resolution of 0.4 (Supplementary Figure 1b, c, Supplementary Tables 2 and 3). The resolution was selected to balance between over-clustering and retaining the ability to identify rare cell types. We then examined the top differentially expressed genes (MAST^24^) for each cluster relative to all other clusters (Figure 2a, b). In two instances, we manually merged two clusters based on the highly overlapping top differentially expressed genes, resulting in a total of 10 cell types. Gene set enrichment analysis was then used to identify hallmark pathways ^25^ enriched for each cell type (Figure 2c). Based on these analyses, we annotated the clusters as nine CAF types and one cluster of pericytes:

**Figure 2.**
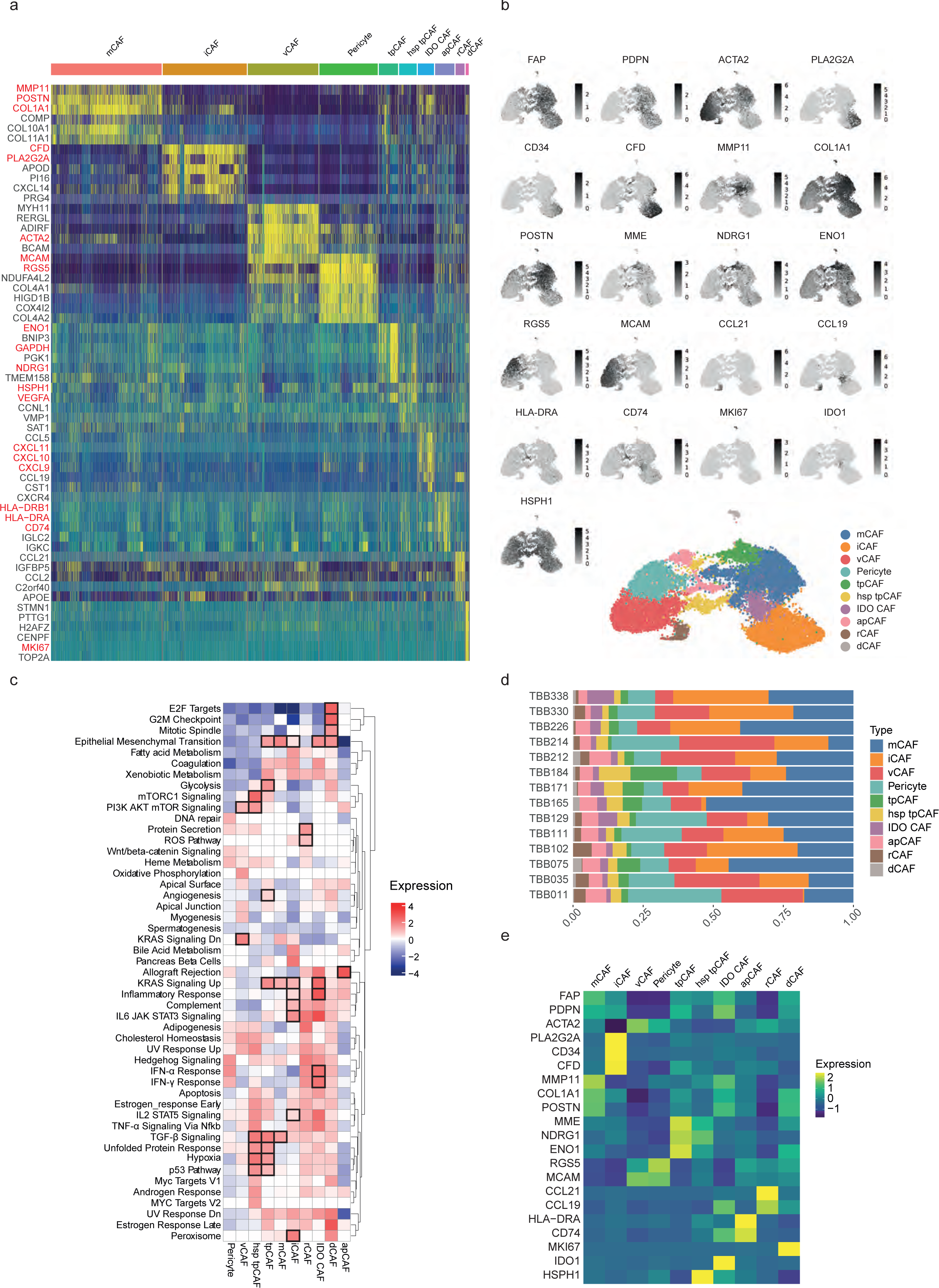
Fibroblast heterogeneity in breast cancer a) Heatmap of the top six differentially expressed genes for each cell type in scRNA-seq data of all stromal cells (n = 16,704). Cellular phenotypes are indicated above the heat map. Key marker genes are highlighted in red. b) UMAPs of all stromal cells coloured by type. Together with a corresponding feature plot showing the expression level of selected marker genes for each cell. c) Gene set enrichment analysis comparing the enrichment of hallmark pathways between CAF types. d) Proportion of all CAF types and pericytes per patient. e) Heatmap showing the average gene expression level of all defined marker genes per identified cell type (after batch correction).

#### Matrix CAFs (mCAFs)

We identified two clusters (clusters 0 and 7), characterized by high levels of expression of genes encoding matrix proteins, in particular matrix metalloproteinase MMP11 and COL1A2 (Figure 2a, Supplementary Figure 1b, Supplementary Tables 2 and 3), based on which we combined these into a single cluster. The resulting group of CAFs (n = 4,525 cells) expressed *MMP11* and other matrix metalloproteinase-encoding mRNAs, collagen-encoding mRNAs (*COL10A1, COL11A1, COL8A1, COL1A2, COL12A1, COL3A1*, *COL8A1*, and *COL5A2*). Further, mRNAs encoding non-collagenous matrix proteins (*COMP* and *POSTN)* were also amongst the top 20 differentially expressed genes of this group (Figure 2 a, b). Cells of this cluster also expressed high levels of genes associated with adhesion (e.g., *LRRC15*, *LRRC17*, and *ASPN*) and with migration (e.g., *POSTN, SULF1*, *INHBA*, and *VCAN*) (Figure 2a, Supplementary Tables 2 and 3). As the top two differentially expressed genes of this cluster were *MMP11* and *POSTN*, we named these cells mCAFs. We conducted a gene set enrichment analysis (GSEA) using hallmark pathways to assess our annotations in an orthogonal manner^25, 26^. For mCAFs, the pathways upregulated included TGF-beta signalling, which is associated with the development of activated myofibroblasts^27^, KRAS signalling, and pathways underlying the epithelial to mesenchymal transition (EMT) (Figure 2c). The EMT-pathway is defined, in this analysis, by multiple collagens, as well as *CTHRC1*, *FAP*, *INHBA*, *LRRC15*, *MGP*, *POSTN*, and *VCAN*. These are all genes associated with matrix remodelling and migration, supporting our mCAF annotation.

#### Inflammatory CAFs (iCAFs)

Cells belonging to the second largest cluster (n = 3,439 cells) were characterized by unique expression of phospholipase encoding *PLA2G2A* and of genes involved in the complement pathway (e.g., *CFD* and *C3)* (Figure 2a, b). In addition, *CD34* was among the top differentially expressed genes in this cluster (Supplementary Tables 2 and 3). CD34 is a marker for hematopoietic stem cells but is known to be expressed by fibroblasts and has previously been linked to inflammation^12, 21^. This cluster also showed high expression of cytokines and chemokines including *CXCL12*, *CXCL14*, and *IL6* (Supplementary Tables 2 and 4). *CXCL12* has previously been used as a marker for CAFs associated with inflammation14,21 and for immune regulatory CAFs^16^, and we thus labelled this cluster iCAFs. Consistent with pro-inflammatory activities, the GSEA revealed that iCAFs are characterized by an upregulation of the IL6-JAK-STAT3 pathway as well as KRAS and complement signalling (Figure 2c). The pancreatic beta cell pathway also showed strong enrichment in iCAFs which is most likely due to a high gene expression level of DPP4 in this cluster (Supplementary tables 2 and 3).

#### Vessel-associated CAFs (vCAFs)

The overall gene expression profile of the CAF cluster with the third highest number of cells (n = 2,886 cells) suggested a link to angiogenesis since these cells showed high expression of *NOTCH3*, an important receptor in vascularisation and angiogenesis^28^, of collagen *COL18A1*, involved in angiogenesis regulation,^29^ and of *MCAM* (which encodes CD146) (Figure 2a, b, Supplementary Tables 2 and 3). These cells did not express the pericyte marker *RGS5* (Figure 2a, b; Supplementary Tables 2 and 3). We named these vCAFs, in line with other descriptions of *RGS5*^-^ fibroblasts characterised in human and mouse cancer^15, 21^. This cluster did, however, show a strong gene expression overlap with *RGS5*^+^ pericytes (e.g., *MCAM* and *ACTA2) (*n=2,389 *RGS5*^+^ pericytes).

#### Tumour-promoting CAFs (tpCAFs)

We observed strong differential expression of proliferation-, migration-, and metastasis-associated genes (e.g., *PDPN*, *MME*, *TMEM158*, and *NDRG1*) as well as stress-response-associated genes (e.g., *ENO1*), and *GAPDH* in a small cluster (n = 786 cells) (Figure 2a, b; Supplementary Tables 2 and 3). This cluster uniquely expressed high levels of *MME* (which encodes CD10), a membrane metalloprotease, and *TMEM158*, an indicator of Ras pathway activation, and also expressed high levels of *VEGFA*, which promotes angiogenesis and vascularisation. Since this gene expression signature resembles that of tumour cells, we named this cluster tpCAFs. This cluster also showed elevated expression of hypoxia marker carbonic anhydrase IX (CA9, Supplementary Tables 2 and 3) suggesting their close proximity to tumour derived hypoxic regions. However, we did not discriminate between hypoxia high and hypoxia low tpCAFs in the scRNA-seq dataset.

A second cluster of tpCAFs (n = 722) expressed high levels of mRNAs encoding heat-shock proteins including HSPH1 and HSP90AA1 (Figure 2a, b, Supplementary Tables 2 and 3) as well as most of the characteristic genes of the first tpCAF cluster. We interpret these cells as tpCAFs that are under higher levels of cellular stress and labelled them heat-shock protein high tpCAFs (hsp_tpCAFs). The GSEA showed an upregulation in both tpCAF clusters of numerous pathways including EMT-associated pathways, TGF-beta and KRAS signalling, as well as glycolysis, MTORC1, and PI3K/Akt/mTOR signalling, the P53 pathway, and hypoxia (Figure 2c). Since many of these upregulated pathways also seen in tumour cells, this is consistent with a role of these clusters of CAFs in promoting tumour growth.

#### IDO CAFs

We annotated a cluster with high differential expression levels of genes associated with chronic inflammation (e.g., *IL32*), as well as of genes upregulated in response to interferons (e.g., *CXCL9*, *CXCL10*, *CXCL11*, and *IDO1*) as IDO^+^ CAFs, due to their strong interferon response (Figure 2a, b; Supplementary Tables 2 and 3). The GSEA of this cluster showed a strong upregulation of the inflammatory response pathways and of interferon-alpha and interferon-gamma responses (Figure 2c), although several additional signalling pathways were enriched in these cells (e.g., IL2-STAT5, TNF-alpha, IL6-JAK-STAT3, and KRAS signalling).

#### Antigen-presenting CAFs (apCAFs)

We identified two clusters (cluster 8 and 10, Supplementary Figure 1b, Supplementary Tables 2 and 3) with high expression of genes involved in MHC-II-associated antigen presentation, including *HLA-DRA*, *HLA-DRB1,* and *CD74* (Figure 2a, b). We manually grouped these clusters based on strongly overlapping top differentially expressed marker genes and named the resulting cluster apCAFs, as previously suggested ^12^. In the GSEA, apCAFs showed the highest upregulation amongst all CAF types of the allograft rejection pathway, in line with their strong expression of MHC-II machinery-related genes (Figure 2c).

#### Reticular-like CAFs (rCAFs)

A relatively rare cluster (n = 373 cells) showed strong differential expression of *CCL21* and *CCL19* (Figure 2a, b, Supplementary Tables 2 and 3). These are markers of reticular fibroblasts in lymphoid tissues that facilitate homing of naïve T cells^30^.

### Dividing CAFs (dCAFs)

The smallest cluster of CAFs (n = 126 cells) in our dataset displayed high expression of genes upregulated during cell division (e.g., *TUBA1B* and *MKI67*, Figure 2a, b; Supplementary Tables 2 and 3). The GSEA supported annotation as dCAFs, showing that E2F targets, the G2M checkpoint, and mitotic spindle pathways were upregulated in cells of this cluster (Figure 2c).

To summarize, differential gene expression analysis together with gene set enrichment analysis allowed us to identify 10 biologically interpretable cell types (9 CAF types and one cluster of pericytes) with unique gene expression profiles. All phenotypes, even rare ones such as rCAFs, were detected in each of our 14 patients (Figure 2d). Further, the cells could broadly be separated into two groups, based on the expression of several genes (Figure 2b, e, Supplementary Figure 1d, Supplementary Table 4). Among others, the two groups could be distinguished by their expression of *FAP* and *SMA*, which have been used in previous studies to define myofibroblasts or CAFs in general and to preselect CAFs before sequencing^27^. We termed the *FAP*^+^/*DCN*^+^ CAFs “activated”; previous work has also referred to these as myofibroblast-like^27^. vCAFs, pericytes, and rCAFs did not show *FAP* expression but did express *ACTA2* (Figure 2e) and we termed them “non-activated”. This is in line with the results from the GSEA which showed upregulation of the KRAS signalling pathway in tpCAFs, mCAFs, iCAFs, IDO CAFs, dCAFs and apCAFs and KRAS downregulation in vCAFs (Figure 2d). Batch correction had no substantial effect on the recovered CAF types (Supplementary Figure 1e).

To enable a straightforward identification of our defined CAF types using different methods, for example flow cytometry or imaging, we sought to define the subset of marker genes that best identified each CAF type. To this end, we examined the top differentially expressed genes for each CAF type and selected those markers that were both in agreement with the current literature ^13, 15, 16, 31^ and also compatible with our goal of conducting a spatial imaging analysis of CAF types within tumour tissue (i.e., proteins with high-quality antibodies for multiplex imaging that should not be confounded with markers for other non-stromal cell types). Our selected markers were *RGS5* for pericytes, *MCAM* for vCAFs, *MME* (together with *NDRG1* and *ENO1*) for tpCAFs, *IDO1* for IDO CAFs, *HSPH1* for hsp_tpCAFs, *MMP11* (together with *COL1A1* and *POSTN*) for mCAFs, *PLA2G2A, CDF* and *CD34* for iCAFs, *MKI67* for dCAFs, *CCL21* (and *CCL19*) for rCAFs, *CD74* and *HLA-DR* for apCAFs, and *PDPN* and *FAP* for the general category of activated CAFs (Figure 2e). When cells from the scRNA-seq dataset were clustered using only these marker genes, we were able to distinguish all our proposed CAF phenotypes (Supplementary Figure 1f). These markers were then used to select markers for subsequent imaging (Supplementary Figure 1g). In summary, we identified nine CAF types and a single group of pericytes in tumours from a cohort of 14 breast cancer patients and identified a set of 17 markers to effectively distinguish between these CAF types.

### A general CAF classification scheme

To test whether our classification scheme is independent of tumour type, we next investigated CAF heterogeneity in four publicly available scRNA-seq datasets from lung cancer^11^, colon cancer^32^, pancreatic ductal carcinoma (PDAC)^33^, and head and neck squamous cell carcinoma cancer^34^. In the lung cancer dataset, we increased the number of fibroblasts by pooling raw data of both the test and validation cohorts from the previous study, resulting in 1,377 CAFs as defined by our criteria and 574 cells from adjacent healthy tissue (Supplementary Figure 2). The colon cancer dataset contained 1,568 CAFs originating from tumour and 1,917 cells from healthy tissue (Supplementary Figure 3). The PDAC dataset contained 1,762 fibroblasts from tumour tissue (Supplementary Figure 4). The head and neck squamous cell carcinoma dataset contained 1,354 cells that we identified as fibroblasts (Supplementary Figure 5); of those, 1,016 fibroblasts originated from tumour tissue, and 315 came from matching lymph node metastases. Taken together, our integrated dataset contained 5,723 CAFs from four different primary tumour types (Figure 3a); we excluded cells from healthy tissue and metastatic sites when integrating the datasets to keep the dataset comparable to our breast cancer dataset and which only included cells from primary tumours. The datasets were integrated using anchors to correct for batch effects as previously described^35^, followed by further batch correction as was performed for the breast cancer dataset (Figure 3a).

**Figure 3.**
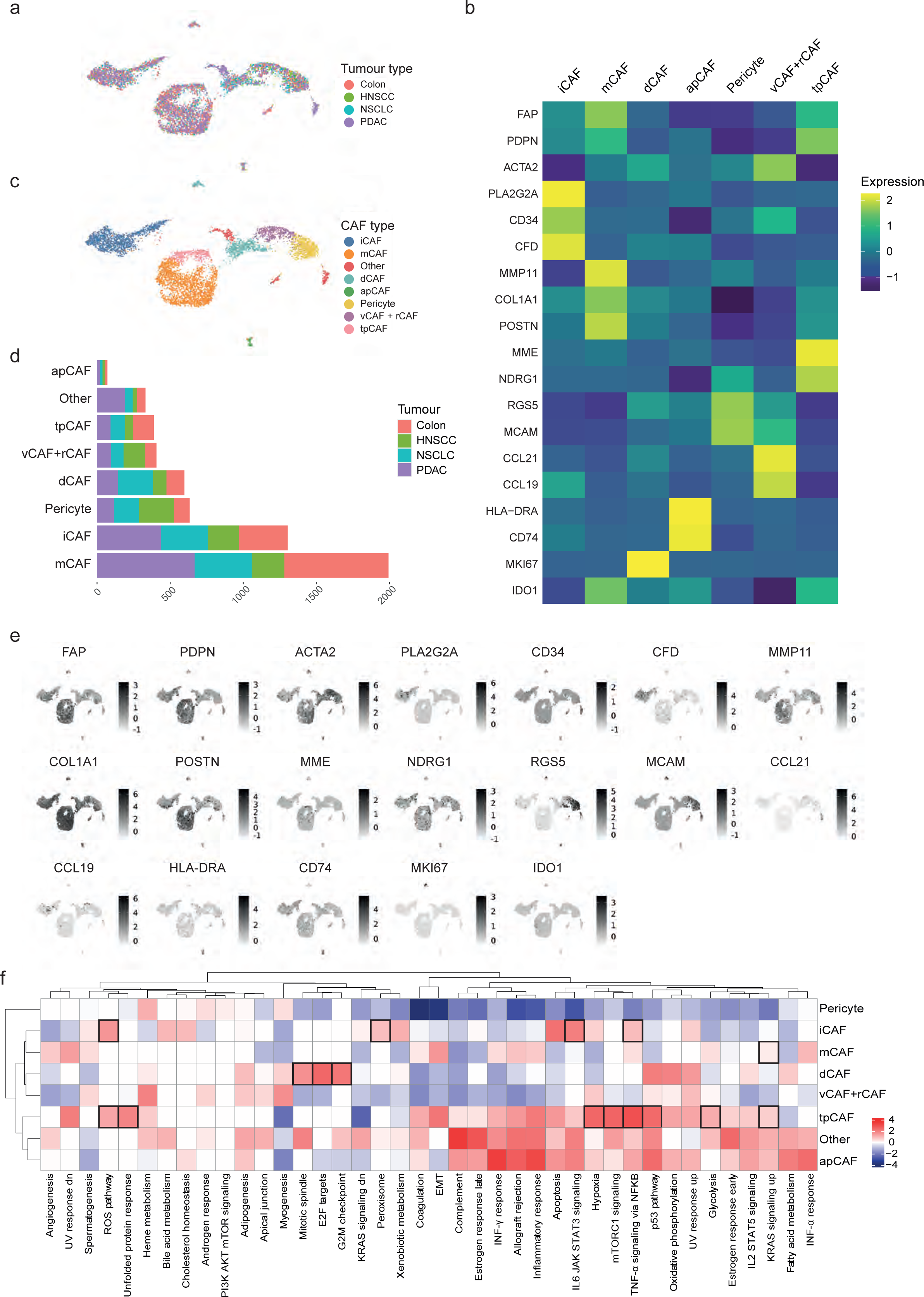
Fibroblast heterogeneity in multiple cancer types a) UMAP showing the validation datasets (lung cancer, head-and-neck squamous cell carcinoma, colorectal cancer, pancreatic ductal carcinoma). b) Heatmap showing the average marker gene expression of each identified cell type in the integrated validation dataset. c) UMAP showing the final CAF classification of the validation cohort. d) Bar chart showing the absolute numbers of all CAF types and pericytes as detected with unbiased clustering of the validation dataset and the respective proportions of each cell type per tumour type. e) Feature plot showing the cellular expression levels of selected marker genes on the UMAP. f) Heatmap showing the results of gene set enrichment analysis for all defined cell types.

We used a two-step approach to identify CAF types in the integrated dataset and to validate our findings. We first performed unbiased clustering on full single-cell gene expression profiles of the integrated dataset and identified all our previously defined CAF types as well as pericytes (Figure 3b, c, Supplementary Figure 6a, b, Supplementary Tables 5 and 6). All CAF types were detected in all cancer types (Figure 3d). Analysis of this integrated dataset yielded similar top differentially expressed genes for each CAF type as identified in the breast cancer dataset (Figure 3b, c, e; Supplementary Tables 5 and 6), and GSEA analysis also showed that similar pathways were upregulated (Figure 3f). KRAS signalling was upregulated in tpCAFs and mCAFs. tpCAFs showed strong expression of the unfolded protein response, hypoxia, mTORC1 signalling, p53 signalling, and glycolysis (pathways Figure 3f). iCAFs showed pathway enrichment of the ROS, peroxisome, IL6/JAK/STAT3 as well as TNF-a signalling pathways (Figure 3f). Clustering the integrated dataset using only our breast cancer-defined CAF marker genes (Figure 3e) also identified all our defined CAF types (Supplementary Figure 6 c-e), providing further support that these markers are sufficient to classify CAFs.

We also clustered each validation dataset individually (Supplementary Figures 2-5). We detected most CAF types in each tumour dataset in these analyses, although dCAFs were typically missing, and vCAFs and pericytes could not be well distinguished in the colon cancer and PDAC datasets. Further, the PDAC dataset was dominated by mCAFs and iCAFs when analysed individually, consistent with the known fibrotic nature of this type of cancer ^12, 20, 33^ Analysis of a larger PDAC dataset (GSE212996) with over 7,000 CAFs showed that, although heavily dominated by mCAFs and iCAFs, tpCAFs, apCAFs and vCAFs were detected (Supplementary Figure 7). Given the high number of cells in the large PDAC dataset compared to the other validation datasets, we did not include it in the integrated analysis as it could potentially drive the clustering results. In our analyses of the individual datasets, we also included cells from healthy adjacent tissue or metastatic lesions to assess the specificity of the CAF types to primary tumour tissue. All CAF types were detected at all tissue sites but in different numbers (Supplementary Figures 2d, 3, 4d). For example, tpCAFs were almost exclusively found in primary tumours (lung: 262 cells in tumour, 11 cells in healthy tissue; colon: 174 cells in tumour, 1 cell in healthy tissue; head and neck squamous cell carcinoma: 284 cells in tumour, 13 cells in metastatic lymph node; two-sided t-test, p = 0.019). In summary, we were able to identify breast cancer-defined CAF types across multiple cancer types, suggesting that we have defined a general CAF classification system and marker genes.

### Spatial distribution of CAF phenotypes in breast tumours

We next used multiplexed IMC^36^ to analyse the spatial distributions of our defined CAF phenotypes in breast cancer samples from the same patients as were analysed with scRNA-seq. The IMC analysis also allowed us to validate our findings at the protein level. We used our scRNA-seq data to guide design of a 41-plex antibody panel to discriminate the different CAF phenotypes (Figure 4a, Supplementary Table 7). We used expression of FAP and PDPN to distinguish myofibroblasts from non-activated, structural CAFs. We used CD146 (encoded by *MCAM*) for identification of vCAFs, CD34 for iCAFs, CD10 (encoded by *MME*) for tpCAFs, CCL21 for rCAFs, and the proliferation marker Ki-67 for dCAFs. Due to the lack of an antibody suitable for detection of MMP11, mCAFs were identified as PDPN^+^/collagen^high^/fibronectin^high^/FAP^+^ cells that did not express CD10, CD34, or CD146. The IMC antibody panel also included markers that allowed identification of tumour cells and immune cells with a focus on T cell subtypes (Figure 4a, Supplementary Figure 8a). Clustering of the scRNA-seq data using the CAF-targeted subset of the 41 markers in this IMC panel showed that we could recover all our defined CAF types (Supplementary Figure 8b).

**Figure 4.**
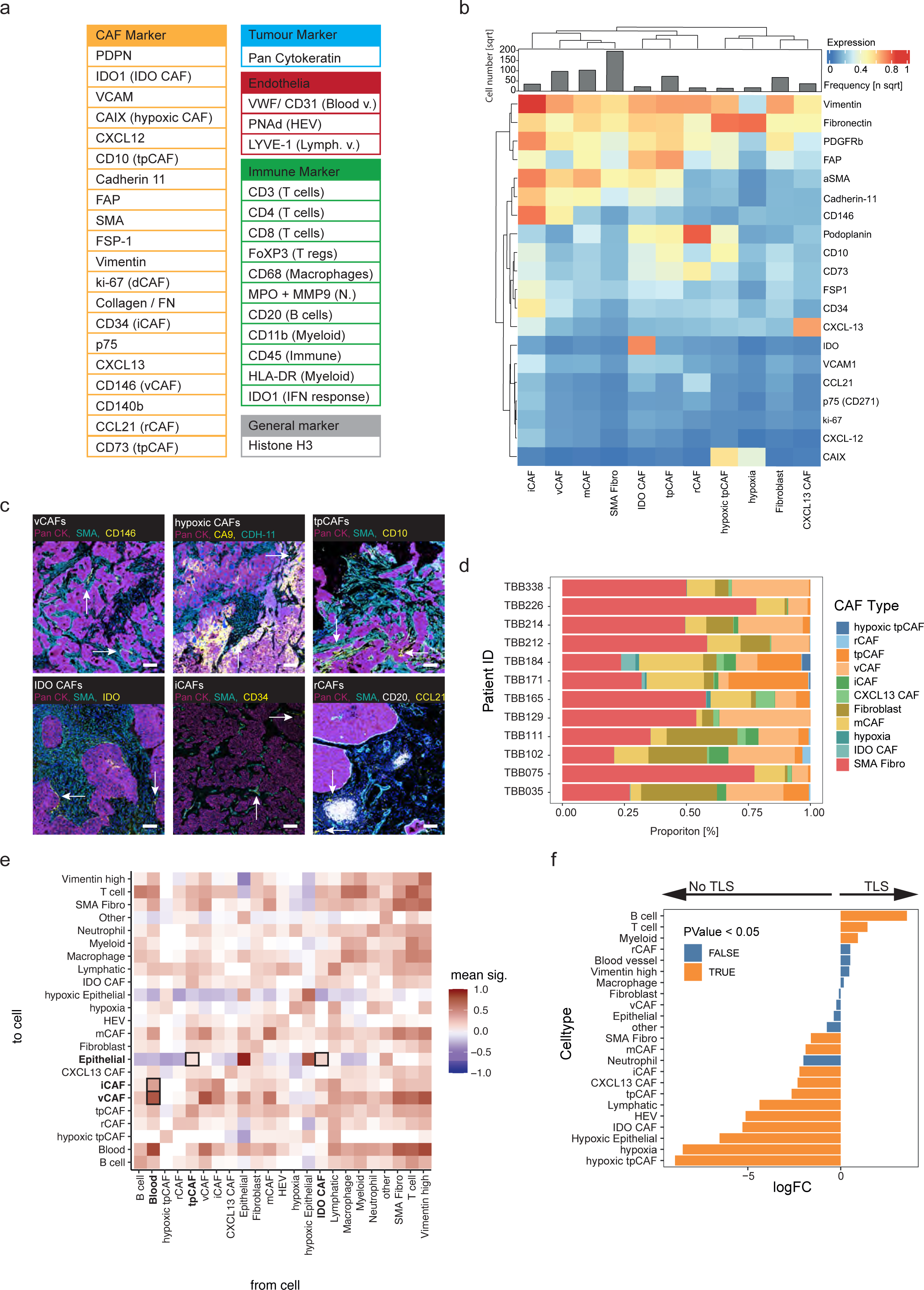
Spatial analysis of CAF types in breast tumours using imaging mass cytometry a) Panel of all markers used in the IMC study. b) Heatmap of marker expression of CAF clusters defined by IMC in breast tumour samples. The histogram indicates the square root of all cell numbers per cluster in each CAF type. c) Images acquired with IMC showing the expression of key markers used in our classification system on the image level. The indicated CAF type is highlighted by arrows. CAFs are identified as follows: vCAFs, CD146; hypoxic CAFs: CDH-11, CA9; tpCAFs, SMA, CD10; IDO CAFs: IDO, SMA; iCAFs: aSMA, CD34; rCAFs: CCL21. PanCK indicates tumour cells, CD20 indicates B cells, Iridium (blue) indicates nuclei in all images. Arrows indicate the respective CAF types in all images. d) Proportion of all CAF types defined by IMC over all patients. e) Neighbourhood analysis showing cell-to-cell interactions at the image level, over all images in the study. The cell-to-cell interactions are compared against a random null-distribution using permutation testing. An interaction score is then generated for each cell-pair based on the p-values calculated on the image level. Positive interaction scores mean that a given pair of cells is neighbouring significantly more often than compared to the null distribution. f) Differential abundance analysis comparing cellular enrichment in TLS containing images versus images not containing any TLS-like structures.

**Figure 5.**
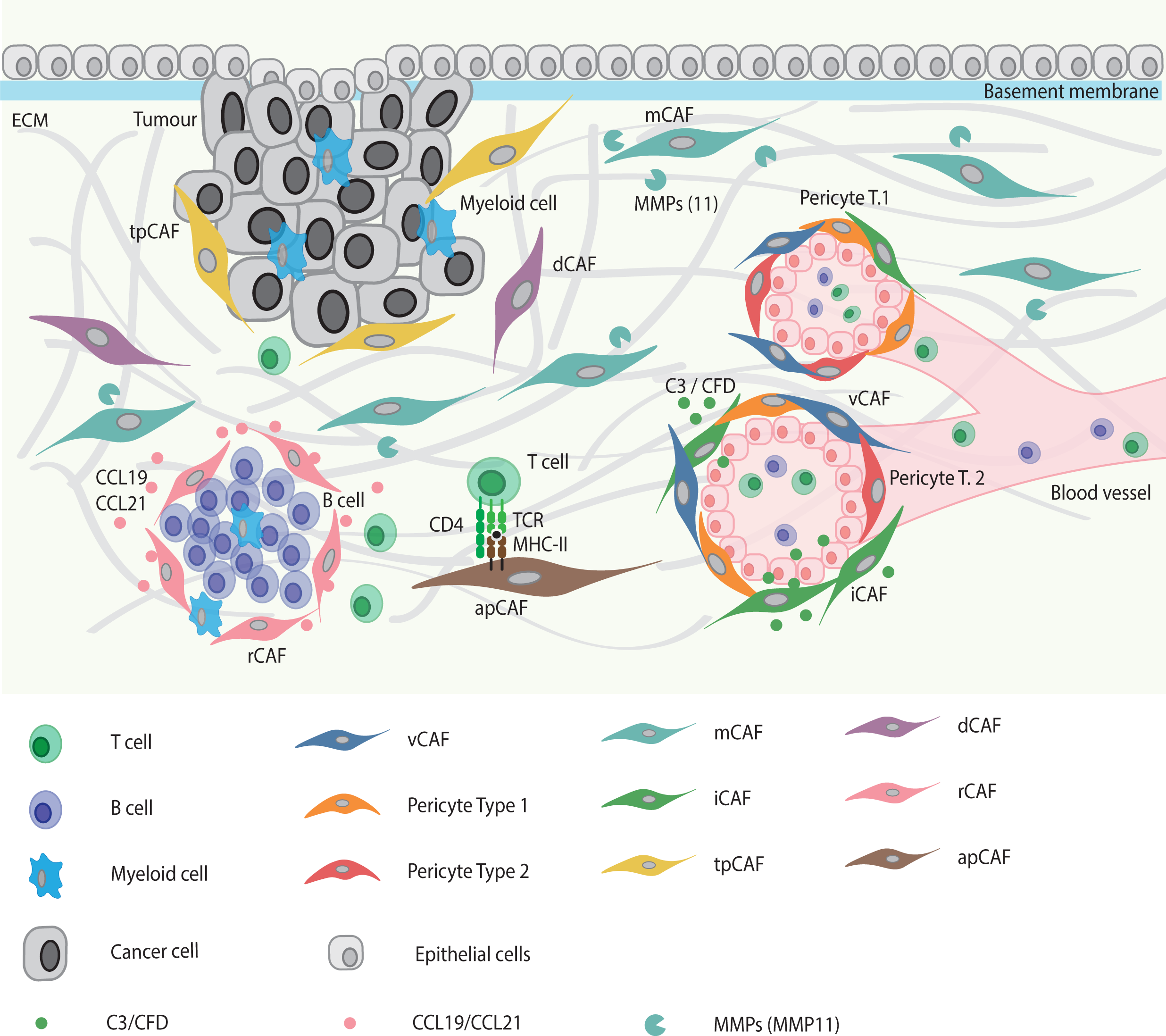
CAF classification scheme Graphical summary showing the spatial distribution patterns of the seven CAF types (excluding apCAFs) in the TME as detected with imaging mass cytometry and their interaction with other cell types.

We stained 12 breast tumour samples (matched tissue samples for 12 of the 14 patient samples analysed by scRNA-seq) with our 41-plex IMC antibody panel and detected stromal, tumour, and immune cells (Supplementary Figure 8c). We selected stroma-rich areas and areas with tertiary lymphoid structures (TLS) based on immunofluorescence imaging (Supplementary Figure 8d), and then analysed these selected areas (7-13 per patient, depending on the number of visible TLS) with IMC. After single-cell segmentation (Supplementary Figure 8e), we identified a total of 222,318 tumour, 104,767 immune, and 140,999 CAFs as well as 29,635 endothelial cells and 55,402 other cells. We identified most of the scRNA-seq-defined CAF subtypes in the IMC dataset (Figure 4b, c). We detected mCAFs, iCAFs, vCAFs, hypoxic and non-hypoxic tpCAFs, IDO CAFs, and rCAFs, but not apCAFs or dCAFs. Most patient samples included multiple CAF types (Figure 4d). Due to the lack of an antibody staining for RGS5, we did not distinguish between vCAFs and pericytes and labelled all CD146^+^/CD31^-^/vWF^-^ cells as vCAFs. Further, since HLD-DR expression is not specific to stromal cells, we did not include it in our analysis and thus did not detect apCAFs.

To study the spatial distributions of CAF types, we conducted a neighbourhood analysis in which we quantified the cell types within a radius of 30 µm of a given cell type, across all imaged regions. vCAFs showed the highest mean interaction score with endothelial cells (Figure 4e) and were exclusively found around endothelial cells in vessel-like structures in the images (Figure 4c). We observed that rCAFs were often found in images with TLS-like structures, where they surrounded aggregated immune cells (mainly CD20^+^ B cells) (Figure 4c) and they further showed a trend towards enrichment in images with TLS structures (Figure 4f). The neighbourhood analysis showed that iCAFs neighbour both vCAFs and endothelial cells (Figure 4c, e). Among CAF types, only CD10^+^/CD73^+^ tpCAFs and IDO CAFs showed positive interaction scores with tumour cells in the neighbourhood analysis, suggesting their close proximity to the tumour (Figure 4e). We investigated this further by comparing the distances of all CAF types to the tumour-stroma border. IDO CAFs were closest to tumour cells with a median distance of 4 µm followed by tpCAFs with a median distance of 14 µm to the tumour-stroma border (Supplementary Fig. 8f), thus confirming that these two CAF types are in relatively close proximity to tumour cells.

In summary, in addition to showing spatial distributions of various CAF types within breast tumours, this IMC analysis has further validated our scRNA-seq-based CAF classification system and showed that most CAF types can be identified in imaging data with only 2-5 markers per type (Table 1). If pericytes are to be distinguished, three additional markers are required. Overall, our demonstration that this classification scheme identifies biologically interpretable CAF phenotypes in multiple cancer types suggests that it will be useful as a general framework for investigations of fibroblast biology.

**Table 1.**
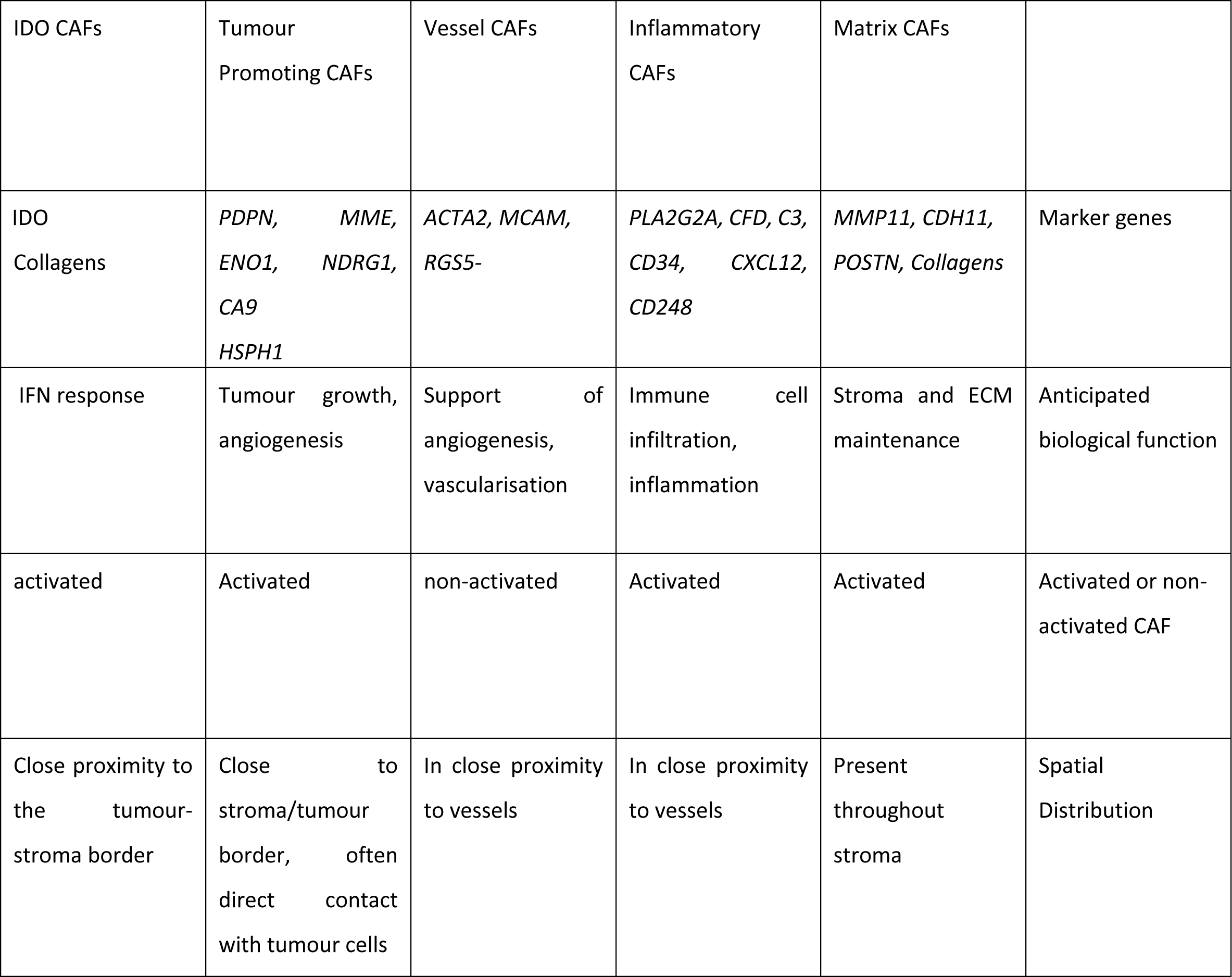
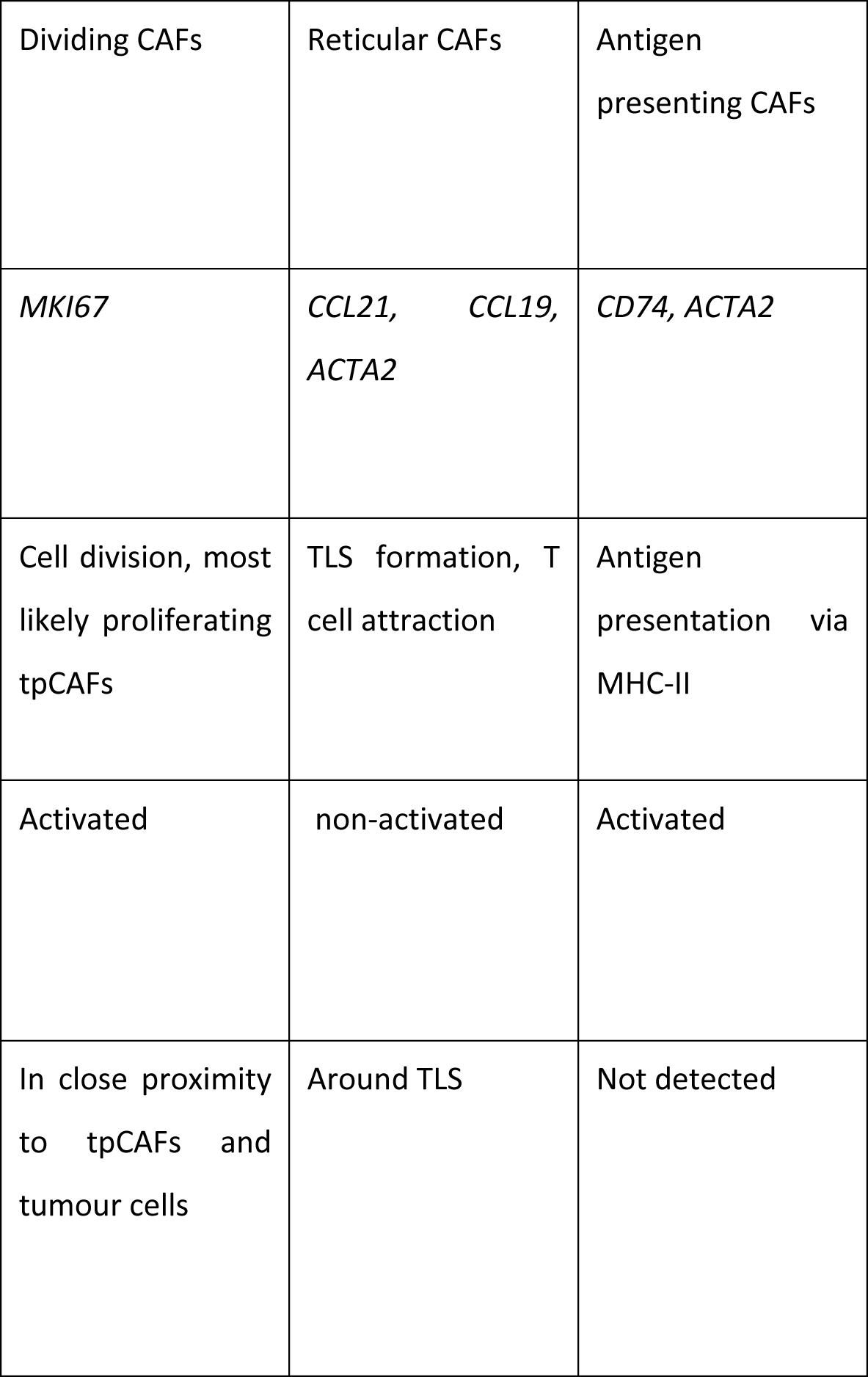
Summary of CAF classification scheme Summarising marker genes, anticipated biological function, classification as either activated or structural CAF, and the spatial distribution for each CAF type as described in this study.

| IDO CAFs | Tumour Promoting CAFs | Vessel CAFs | Inflammatory CAFs | Matrix CAFs |  |
| --- | --- | --- | --- | --- | --- |
| IDO<br>Collagens | *PDPN*,<br>*ENO1*,<br>*CA9*<br>*HSPH1* | *ACTA2*, *MCAM*,<br>*RGS5*- | *PLA2G2A*, *CFD*, *C3*,<br>*CD34*, *CXCL12*,<br>*CD248* | *MMP11*, *CDH11*,<br>*POSTN*, *Collagens* | Marker genes |
| IFN response | Tumour growth,<br>angiogenesis | Support of<br>angiogenesis,<br>vascularisation | Immune cell<br>infiltration,<br>inflammation | Stroma and ECM<br>maintenance | Anticipated<br>biological function |
| activated | Activated | non-activated | Activated | Activated | Activated or non-activated CAF |
| Close proximity to<br>the tumour-stroma border | Close to<br>stroma/tumour<br>border, often<br>direct contact<br>with tumour cells | In close proximity<br>to vessels | In close proximity<br>to vessels | Present<br>throughout<br>stroma | Spatial<br>Distribution |

| Dividing CAFs | Reticular CAFs | Antigen presenting CAFs |
| --- | --- | --- |
| *MKI67* | *CCL21*, *CCL19*,<br>*ACTA2* | *CD74*, *ACTA2* |
| Cell division, most likely proliferating tpCAFs | TLS formation, T cell attraction | Antigen presentation via MHC-II |
| Activated | non-activated | Activated |
| In close proximity to tpCAFs and tumour cells | Around TLS | Not detected |

## Discussion

A universal a classification system for CAFs has been lacking^10^, although studies of CAF heterogeneity in many cancer types have led to definition of numerous CAF subtypes^12–14, 16, 21^. Previous studies were often limited regarding their sample size or marker availability or used single markers to define cell phenotypes. Thus, rarer cell types were not always detected^21^, and most CAF phenotypes have been defined in single cancer types. In addition, the spatial distributions of CAF phenotypes in tissue have not been previously investigated in detail.

Based on our analysis of scRNA-seq data of more than 16,000 stromal cells from 14 human breast cancer tumour samples, coupled with multiplex single-cell imaging of matched samples, we identified marker expression and spatial distribution of nine CAF types as well as pericytes. We could further identify these phenotypes in four additional cancer types based on previously published scRNA-Seq data, suggesting that these phenotypes and the classification system we have defined is generalizable.

Our identified phenotypes share markers with previously defined CAF phenotypes^12, 15, 16, 21^ especially those previously found in mouse models of breast cancer^15, 16^. Cells we identified as vCAFs based on expression of *MCAM*, *NOTCH3*, and *COL18A1* are analogous to the vascular CAFs defined in mouse^15^, and the cells we define as dCAFs are analogous to the cycling CAFs identified in mouse^15^. Inflammatory CAFs and immune regulatory CAFs were previously defined based on expression of *Il6*, *Cxcl12*, and *Cxcl1* and *IL6* and *CXCL12*, respectively, in scRNA-seq data from mouse cancer models^7, 16^ and human cancer samples^7, 14, 16, 21, 37^. Although these patterns were recapitulated in our data, we defined a single category of *PLA2G2A^+^*/*CD34^+^*/*CFD*^+^ iCAFs, since all the cells contained within this phenotype expressed genes actively involved in shaping the immune environment. Friedman et al. described ECM-CAFs based on expression of *Fbn1*, which encodes fibronectin^16^. *Fbn1* is highly expressed by our mCAFs, but we found that *MMP11,* also used by Wu et al. to describe myofibroblast-like CAFs^21^, was more strongly expressed in mCAFs. Finally, our apCAFs, defined by their expression of mRNAs encoding MHC-II molecules, were previously described by Elyada et al. in human and mouse PDAC^12^.

Pericytes in human breast cancer, defined by the expression of *RGS5*, have previously been shown to have gene expression patterns similar to those of CAFs^21^. Wu et al. also defined immature and developed perivascular-like cells^21^, offering a potential trajectory from pericytes to vCAFs. Our vCAFs share features with both pericyte types but are more closely related to the developed perivascular-like cells by Wu et al.^21^. It remains to be determined whether pericytes are an independent cell type or whether they are a subset of specialised fibroblasts found in both healthy and diseased tissues.

rCAFs are closely related to vCAFs based on the UMAP. They express *CCL21* and *CCL19* and as such, are similar to reticular fibroblast cells found in lymphoid structures. *CCL19*-expressing CAFs have been previously described in cancer, usually as part of larger CAF groups^14, 38^. We chose to define rCAFs as an individual category, despite their relative rarity, since we detected them mostly in close proximity to TLS in the IMC analysis.

The fact that we were able to define almost all CAF phenotypes in datasets from multiple cancer types, both when analysed individually and when analysed as an integrated dataset, strongly suggests that CAF phenotypes are generalizable across cancer types. A recent pan-cancer study has compared plasticity of fibroblasts and showed that distinct features are shared between the different tumour sites^39^.This may extend also to subtypes; for example, our breast cancer dataset mainly consisted of luminal B cancer, but the CAF phenotypes we defined were also present in samples of the other subtypes. Still, our data also indicate that different cancer types may be dominated by different CAF types. In the case of highly fibrotic PDAC, for instance, we were able to identify all CAF types but the main clusters were mCAFs and iCAFs.

Further, it is likely that the CAF phenotypes we have identified represent a fluid spectrum of phenotypes with different functional features rather than fixed cellular states. Consistent with this, several of the phenotypes cluster closely together in the UMAP space, and clustering with subsets of markers rather than the entire measured transcriptome show small fluctuations in the distribution of CAF phenotypes. Time course experiments in model systems would be needed to study such relationships further. In addition, although we have defined each CAF phenotype based on differentially expressed genes that suggest a dominant biological feature or function, these cell types may very well have more than one biological function. For example, although we defined mCAFs based on expression of genes involved in matrix remodelling and production, they can also produce inflammatory factors such as cytokines and chemokines and some express genes known to facilitate adhesion and migration without it being their defining feature. Similarly, while tpCAFs mainly express genes indicative of tumour mimicking, they also express MMPs and matrix proteins and may therefore also function to remodel the ECM.

We provide here a general CAF classification system and a 17-marker panel that we have validated across cancer types; this panel differentiates between activated and non-activated CAFs, and identifies the functional categories of vessel CAFs, matrix CAFs, immune CAFs, tumour-promoting CAFs, IDO CAFs, antigen-presenting CAFs, dividing CAFs, and reticular-like CAFs. Should the nature of follow up studies require different markers (e.g. investigation of a particular functional subpopulation of CAFs), our dataset can serve as a reference for choosing the most appropriate marker genes. In either context, our concepts and dataset should establish a basis for future studies of this important cell type.

## Supporting information

Supplementary Table 1_ Clinical Data and 10X Run Information

Supplementary Table 2_ Breast cancer clusters res04

Supplementary Table 3_ Breast cancer CAF types

Supplementary Table 4_ Breast cancer CAF activation state

Supplementary Table 5_Integrated Data clusters res08

Supplementary Table 6_Integrated Data CAF types

Supplementary Table 7_ Antibody Panel

## Abbreviations

CAFs: cancer associated fibroblasts
ECM: extracellular matrix
IMC: imaging mass cytometry
MMPs: matrix metalloproteinases
scRNA: seq single cell RNA sequencing
TLS: tertiary lymphoid tissue
TME: tumour microenvironment
vCAFs: vascular CAFs
mCAFs: matrix CAFs
iCAFs: inflammatory CAFs
tpCAFs: tumour promoting CAFs
hsp_tpCAFs: heat shock protein high tpCAFs
rCAFs: reticular-like CAFs
dCAFs: dividing CAFs
apCAFs: antigen presenting CAFs

## Author Contributions

L.C. generated the IMC dataset, analysed all scRNA-seq and IMC data, and wrote the manuscript. S.T. generated the scRNA-seq dataset and provided input on the manuscript and analysis. T.A., C.L., and M.R. provided the samples as well as clinical input. N.dS. provided critical input on the manuscript. B.B. supervised the work and provided input at all levels.

## Materials and Methods

### Lead contact

All questions regarding the manuscript should be directed to lead contact Bernd Bodenmiller.

### Materials availability

There were no new reagents generated in this study.

### Data and code availability

All data and analysis code will be publicly available after publication. The breast cancer RNA sequencing data have been added under with the accession number E-MTAB-10607 to the ArrayExpress database at EMBL-EBI (www.ebi.ac.uk/arrayexpress). Publicly available datasets used in this study are available at the Gene Expression Omnibus with the following accession numbers: GSE132465 (colon cancer), GSE1547 PDAC (small), GSE21 PDAC (big), GSE103322 (head and neck squamous cell carcinoma) and in ArrayExpress at EMBL-EBI under the accession numbers E-MTAB-6149 and E-MTAB-6653 (lung cancer).

### Code availability

All analysis code will be made publicly available upon publication.

### Clinical samples

All primary breast cancer samples included in this study were previously collected in collaboration with the Patient’s Tumor Bank of Hope (PATH, Germany) and analysed by CyTOF^40^, scRNA-seq, and IMC (Tietscher et al., in press) for in-depth characterization of tumour and immune landscapes. All tissue and health-related data were collected under approval of the Ethics Committee Zurich (#2016-00215) and the faculty of medicine ethics committee at Friedrich-Wilhelms-University Bonn (#255/06). The fibroblast scRNA-seq data analysed in this study are part of a dataset previously generated by our laboratory (Tietscher et al., in press). The presented IMC data were specifically acquired for this study. For two of the 14 breast cancer samples for which scRNA-seq data were available, IMC measurements were not possible due to missing patient consent for FFPE-based analysis or due to low-quality FFPE material. Tumour subtypes in this study were defined as follows: Luminal A (ER^+^ and/or PR^+^, HER2^-^, Ki-67^+^ < 20%), Luminal B (ER^+^ and/or PR^+^, HER2^-^, Ki-67 ≥ 20%), Luminal B-HER2^+^ (ER^+^ and/or PR^+^, HER2^+^), HER2^+^ (ER^-^, PR^-^, and HER2^+^), and triple negative (ER^-^, PR^-^, and HER2^-^).

### Breast cancer – scRNA-seq dataset and fibroblast identification

The scRNA-seq dataset presented here was previously generated using the 10x Genomics platform, and the raw data pre-processing, quality control steps, and main cell type annotation have been described (Tietscher et al., submitted). Briefly, gene-by-cell matrices were generated from the raw sequencing data using CellRanger (10x Genomics, v3.0.1) and subsequently transformed into Seurat objects (Seurat v3.0.2). After removing high-confidence doublets using the DoubletFinder Package, all Seurat objects were merged, and single cells with >7500 or <200 genes, with >75000 read counts, or with >20% of reads mapping to mitochondrial RNA were excluded. Highly variable genes were identified by the *sctransform* wrapper in Seurat and used to construct principal components. The principal components covering the highest variance in the dataset were used as input for graph-based clustering. Differential gene expression analysis was performed for the resulting clusters, and main cell types were annotated based on the cluster expression of established marker genes (*EPCAM* and *CDH1* for epithelial cells, *PECAM1* and *VWF* for endothelial cells, *PDGFRB* and *FAP* for fibroblasts, *CD3*, *CD4*, *CD8*, and *NCR1* for the T and NK cell fraction, *CD14*, *ITGAX*, and *HLA-DRA* for myeloid cells, *MS4A2* for mast cells and basophils, *MS4A1* for B cells, and immunoglobulin-encoding genes for plasma cells). For the present study, all clusters annotated as “fibroblasts” in the original dataset were used for downstream analysis.

### Breast cancer – analysis of fibroblast clusters

We used the Seurat package (v4.0.2) function *sctransform* to normalise and scale the data, using the variables “percent.mt” (mitochondrial genes), “percent.krt”, and “percent.MGP” for regression (*KRT* and *MGP* were detected across all cell types in some samples due to contamination originating from apoptotic tumour cells). We used the Seurat harmony batch correction wrappers to reduce a visible sample effect. After running principal component analysis, the first 25 components were used for both graph-based clustering using the Seurat functions *FindNeighbours* and *FindClusters* as well as dimension reduction analysis such as UMAP. We used different resolutions in resolution steps of 0.1 for clustering and investigated the clustering hierarchy by plotting a clustree^41^. Use of results of clustering at resolution 0.4 resulted in a total of 12 clusters (numbered 0 through 11, Supplementary Figure 2c, Supplementary Table 4). Differential gene expression per cluster was analysed using the *FindAllMarkers* function using MAST testing (min.pct = 0.25, logfc.threshold = 0.25). Due to similar gene expression patterns, clusters 0 and 7 were unified as mCAFs, and clusters 8 and 10 were unified as apCAFs. Cluster 1 was assigned as iCAFs, cluster 2 as vCAFs, cluster 3 as pericytes, cluster 4 as tpCAFs, cluster 5 as heat-shock protein-high tpCAFs, cluster 6 as IDO CAFs, cluster 9 as rCAFs, and cluster 11 as dCAFs.

### Gene set enrichment analysis

We used the singleseq package^26^ to run gene set enrichment analysis and compare the enrichment of hallmark pathways ^25^ between our defined CAF types.

### Analysis of validation datasets

The head and neck squamous cell carcinoma, colon cancer, and PDAC datasets were downloaded from the NCBI Gene Expression omnibus (GSE103322, GSE132465, GSE154778 and GSE212966, respectively). The fibroblast subsets were then scaled and clustered as described above. For the **head and neck squamous cell carcinoma** dataset, the original fibroblast identification was used. They were then clustered and the resolution of 0.6 was used for fibroblasts resulting in 10 clusters (0, vCAF; 1, vCAF; 2, iCAF; 3, tpCAF; 4, Pericyte; 5, rCAF + apCAFs; 6, tpCAF; 7, Pericyte; 8, other). The **colon cancer** dataset was clustered and cluster 6 at the resolution of 0.2 was identified as fibroblasts. Fibroblasts were then clustered at resolution 0.3 obtaining nine clusters (0, iCAF; 1, mCAF; 2, iCAF; 3, mCAF; 4, Pericyte; 5, mCAF; 6, other; 7, tpCAF; 8, rCAF). Cells identified as “other” were excluded from subsequent analysis. For the **lung cancer** dataset, RAW sequencing files (E-MTAB-6149, E-MTAB-6653) were downloaded and analysed using the CellRanger pipeline (v6.0.0, refdata-gex-GRCh38-2020-A) to obtain gene-by-cell matrices before being analysed in a Seurat object. Fibroblasts were identified from both datasets (E-MTAB-6149, cluster 9 at resolution 0.2; E-MTAB-6653, cluster 12 at resolution 0.2) and merged into one Seurat object. The final clustering of the lung cancer fibroblasts was obtained at the resolution of 0.8 to yield 15 clusters (0, mCAF; 1, tpCAF; 2, mCAF; 3, rCAF; 4, iCAF; 5, other; 6, apCAF; 7, iCAF; 8, pericyte; 9, iCAF; 10, epithelial; 11, other; 12, vCAF; 13, iCAF; 14, mitochondrial count high). The small PDAC dataset (GSE154778) all cells were clustered, and fibroblasts identified in clusters 1 and 6 at resolution 0.1 Fibroblasts were then clustered and resolution of 0.4 was chosen for the final cluster assignment resulting in a total of 5 clusters (0, iCAF; 1, mCAF; 2, apCAF; 3, pericytes; 4, other). For the bigger PDAC dataset (GSE212996) all cells were clustered, and fibroblasts identified as clusters 2 and 4 at the resolution of 0.1. For fibroblasts, a clustering resolution of 0.6 was chosen, resulting in a total of 10 clusters (0, mCAF; 1, mCAF; 2, iCAF; 3, tpCAF; 4, vCAF; 5, apCAF; 6, other; 7, pericyte; 8, iCAF; 9, other). All cells identified as other, epithelial, or mitochondrial high were excluded from subsequent analyses. The different clustering resolutions resulted from the varying sizes of the datasets and our aim to also detect the smaller subclusters.

### Dataset integration

All scRNA-seq datasets were integrated using the in-built *SelectIntegrationFeatures and FindIntegegrationAnchors*functions from the Seurat package. Batch correction was run using the harmony wrapper function for Seurat. Principle component analysis was performed, and the first 15 components were used in UMAP analysis. The integrated fibroblasts were analysed according to the previously described pipeline. Clustering at the resolution of 0.8 resulted in 12 clusters (0, iCAF; 1, mCAF; 2, mCAF; 3, pericyte; 4, dCAF; 5, vCAF + rCAF; 6, mCAF; 7, tpCAF; 8, iCAF; 9, other; 10, other; 11, dCAF; 12, apCAF; 13, other). When clustering the dataset using only the selected marker genes, this resulted in a total of 16 clusters at resolution 0.8 (0, iCAF; 1, mCAF; 2, mCAF; 3, rCAF + pericyte; 4, pericyte; 5, tpCAF; 6, IDO_CAF; 7, iCAF; 8, pericyte; 9, tpCAF; 10, iCAF; 11, rCAF + pericyte; 12, iCAF; 13, iCAF; 14, dCAF; 15, apCAF).

### Tissue preparation and staining

Using a series of HistoClear and graded ethanol solutions, FFPE tissue sections were deparaffinised and rehydrated before antigen-retrieval in a decloaking chamber for 30 minutes at 95 °C using HIER buffer (pH 9.2). Tissues were blocked with 3% BSA for 45 minutes before being stained with the first metal-tagged antibody (anti-SMA, Supplementary Table 7) at room temperature for 4 hours. After washing, the tissue was incubated with a fluorescently labelled anti-mouse (Abcam, AF 555, goat anti-mouse (H+L), catalogue number A21422; RRID_AB_2535844, polyclonal) for 1 hour at room temperature before another washing step and final incubation with Hoechst (dilution 1:500) for 5 minutes. Afterwards, tissue was incubated with the remaining metal-tagged antibodies (Supplementary Table 7) at 4 °C overnight. After washing the next day, the samples were stained with an iridium DNA intercalator before being dried using pressured air.

### Immunofluorescence imaging

Whole slide scans were preformed using a Zeiss AxioScan.Z1 with x10 magnification. The resulting images were used to select between 6-8 stroma rich areas, and, if present, TLS regions, of 1 mm^2^ per patient for IMC imaging analysis (Supplementary Figure 8d).

### Imaging mass cytometry

Brightfield scans of the slides were performed to detect the areas of interest to be measured by IMC. For subsequent imaging, the Hyperion Imaging System, coupled to a Helios time-of-flight mass cytometer was used at a laser intensity of 400 Hz (resolution at 1 µm). A compensation slide was also analysed to account for potential spill-over between the metals. The machine was tuned daily to account for machine performance variability. All markers were checked for their quality of staining (Supplementary Figure 9).

### Analysis

Using the lab’s analysis pipeline (github.com/BodenmillerGroup/ImcSegmentationPipeline), tiff files were generated from the raw data. Images were produced using histoCAT^42^. The tiff files were used for cellular segmentation based on nuclear and membrane markers using the ilastik software^43^. First, fibroblasts were reduced to their nuclear core to reduce cellular overlap and signal falsification (Supplementary Figure 9). *CellProfiler* (v3.1.9)^44^ was then used to generate cell masks and to calculate mean intensities of each marker per cell. Neighbours within a radius of 30 µm of each cell were also identified by *CellProfiler*. The single-cell data were analysed using R (v4.0.4); raw counts were censored excluding the 99.99 percentile before being arc-sinh transformed using cofactor 1.

Graph-based clustering was carried out using the PhenoGraph clustering algorithm. In the first clustering step, the dataset was divided into the three compartments, tumour, immune, and stromal cells (Supplementary Figure 8a, c). Cells identified as immune or stromal were then clustered individually resulting in immune and stromal cell subtypes). CAFs were clustered using *FLOWSOM* clustering as integrated in the *Catalyst* R package ^45^. We excluded HLA-DR from the CAF clustering as it resulted in too many false positive CAFs. This is due to the nature of HLA-DR expression on myeloid cells which are often wide-spread in the stroma and can thus cause false positive signal on other cell types. This was the only marker for which we observed this false positive signal. The neighbourhood analysis was carried out analysing the 15 nearest cells in a defined radius of 25 µm of each cells’ centroid using *imcRtools* (Bioconductor dpi).

### Image generation

Images shown in this study were generated using *histoCAT*-web and *cytomapper* (Bioconductor).

## Supplementary figure titles and legends

**Supplementary Figure 1.**
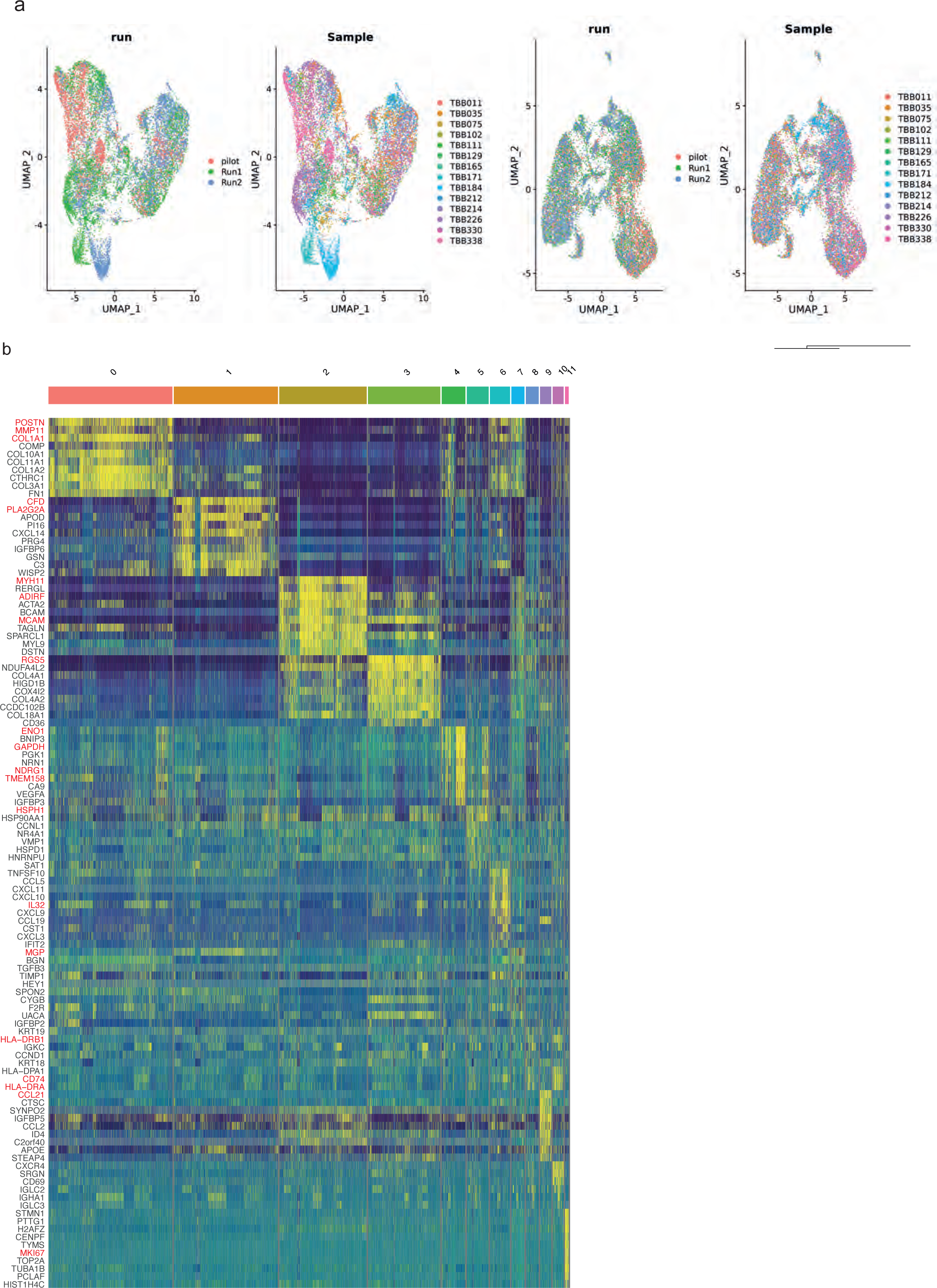
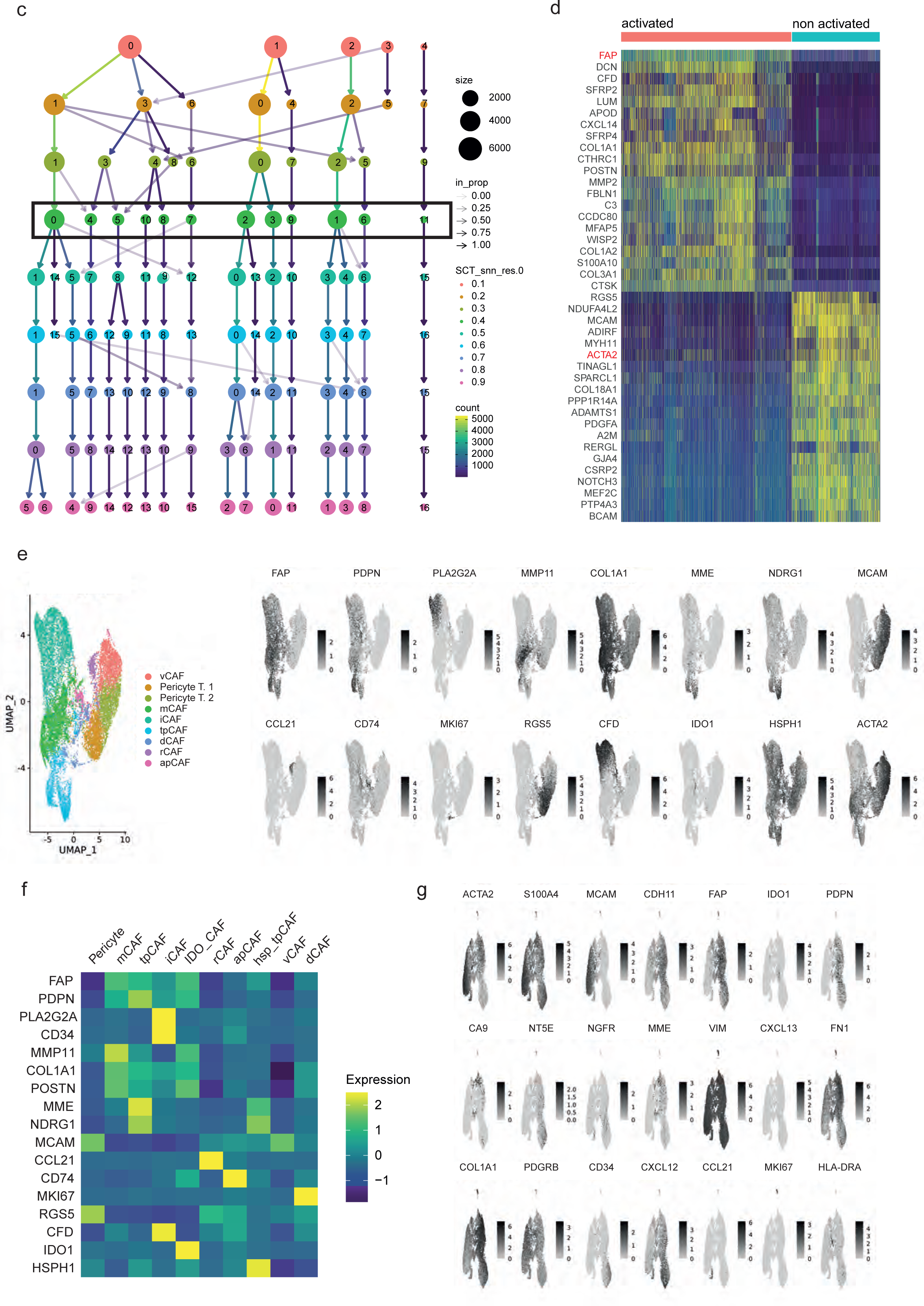
scRNA-seq data of breast cancer a) UMAPs of the uncorrected raw data (left two plots) and the batch corrected data (right two plots), coloured by run and sample ID. b) Heatmap showing the top 10 differentially expressed genes for each cluster at the resolution of 0.4. c) Clustree showing hierarchical clustering for breast cancer fibroblasts. d) Heatmap of all cells showing the classification as activated and non-activated and their respective differentially expressed genes. e) UMAP of CAF type assignment from raw, uncorrected scRNA-Seq data. The feature plots show expression level of our defined CAF selected marker genes. f) Heatmap showing the average marker gene expression of all defined marker genes for clusters detected when clustering all stromal cells using only the subset of defined marker genes. g) Feature plot showing expression of all CAF markers selected for IMC as detected in the breast cancer cohort.

**Supplementary Figure 2.**
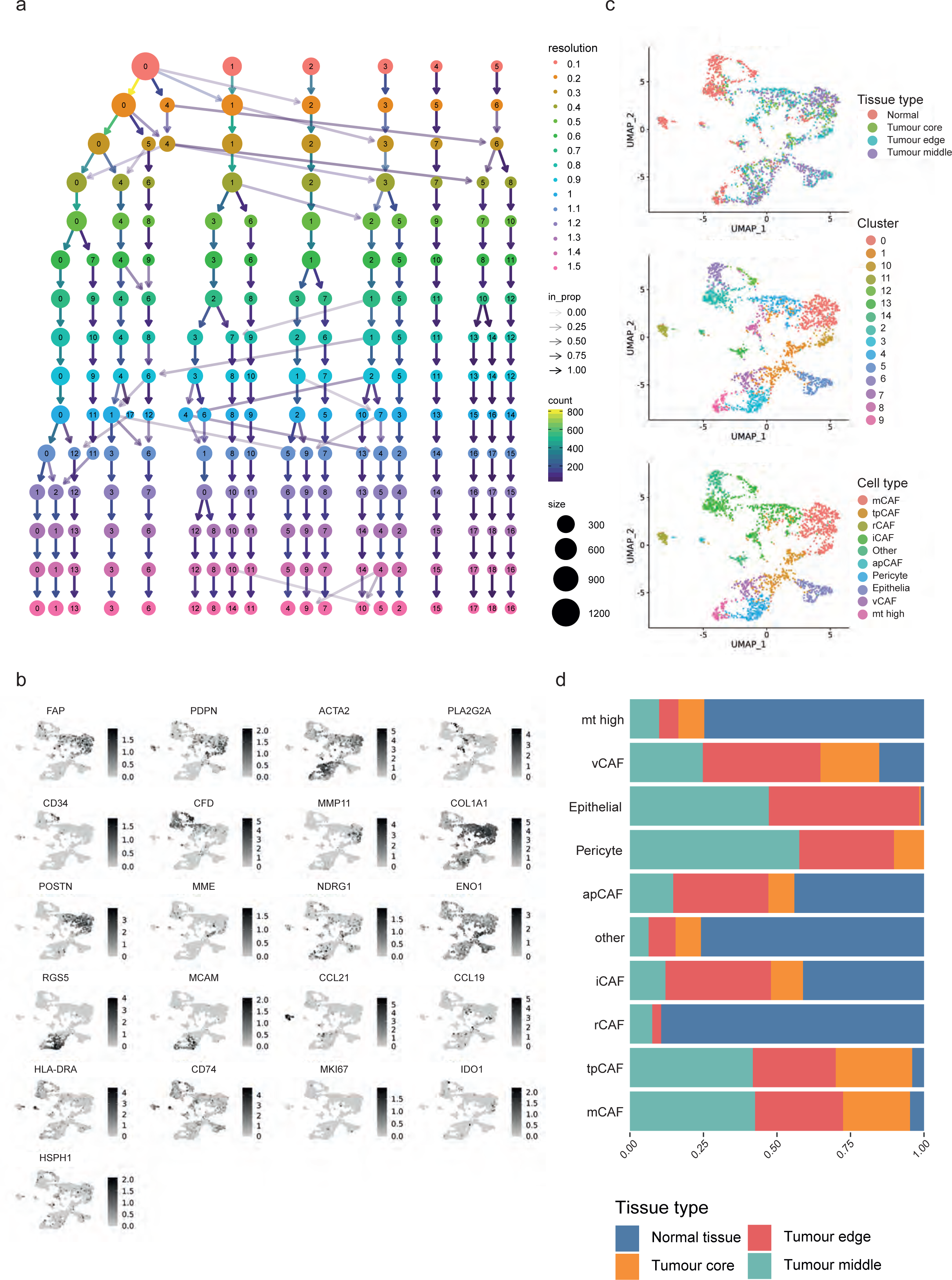
Analysis of the lung cancer scRNA-seq data a) Clustree showing the hierarchical clustering of all lung cancer stromal cells. b) Feature plot showing the marker expression of all marker genes for all cells in this dataset. c) UMAPs coloured by (top to bottom) tissue sites sequenced in the original study (normal tissue, tumour core, tumour edge, tumour middle), clusters obtained at clustering resolution of 0.8, and the final cell type assignments of mCAFs, tpCAFs, rCAFs, iCAFs, other, apCAFs, pericytes, epithelial cells, vCAFs, and mitochondrial-high cells (mt high). d) Proportions of tissue site distribution of all defined cell types in this dataset.

**Supplementary Figure 3.**
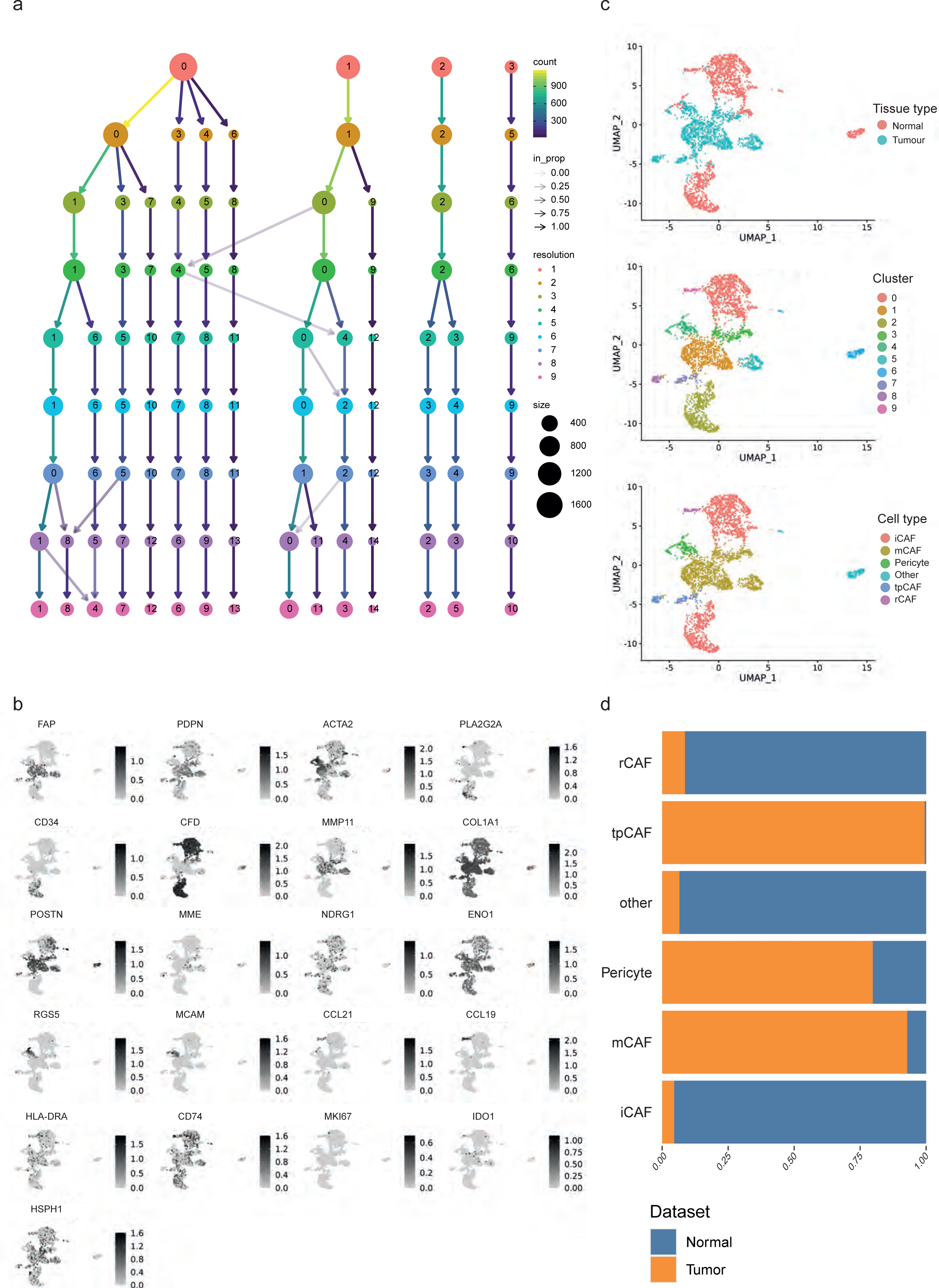
Analysis of scRNA-seq data from colon cancer a) Clustree showing the hierarchical clustering of all colon cancer stromal cells. b) Feature plot of all colon cancer CAFs showing the expression of defined marker genes. c) UMAPs coloured by (top to bottom) tissue sites sequenced in the original study (normal tissue, tumour tissue), clusters obtained at clustering resolution of 0.3, and final cell type assignments of iCAFs, mCAFs, pericytes, other, tpCAFs, and rCAFs. d) Cell type distribution between tumour and healthy tissue origin for all cell types defined in this dataset.

**Supplementary Figure 4.**
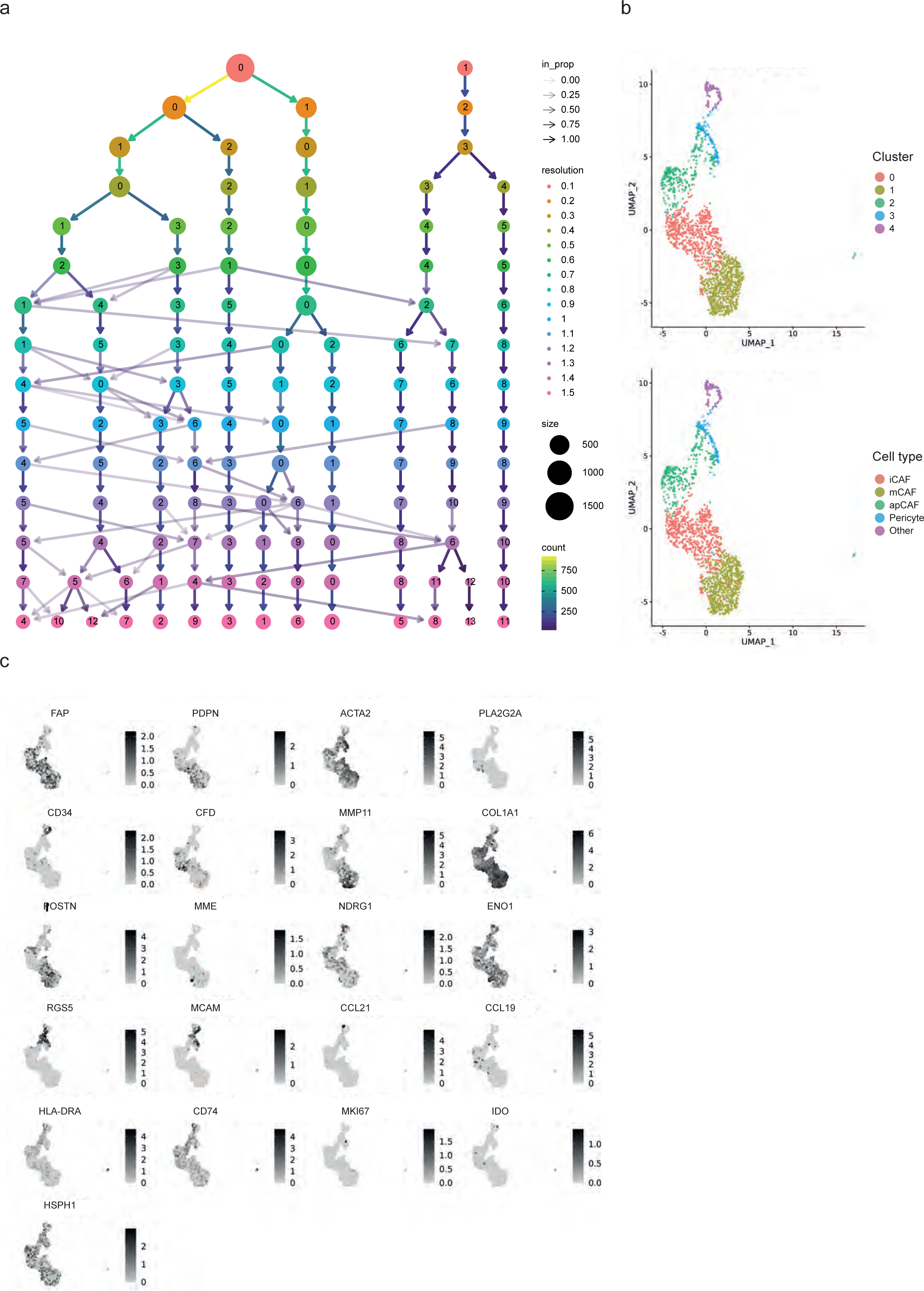
Individual analysis of the small PDAC dataset. a) Hierarchical clustering tree (using the clustree package) showing the clustering results of the small PDAC dataset at clustering steps. b) UMAPs coloured by the chosen clustering resolution of 0.4 and the resulting 5 different clusters as well as the final cluster assignment (iCAFs, mCAFs, apCAFs, pericytes and other). c) Feature plot showing the expression of all previously identified marker genes on the UMAP.

**Supplementary Figure 5.**
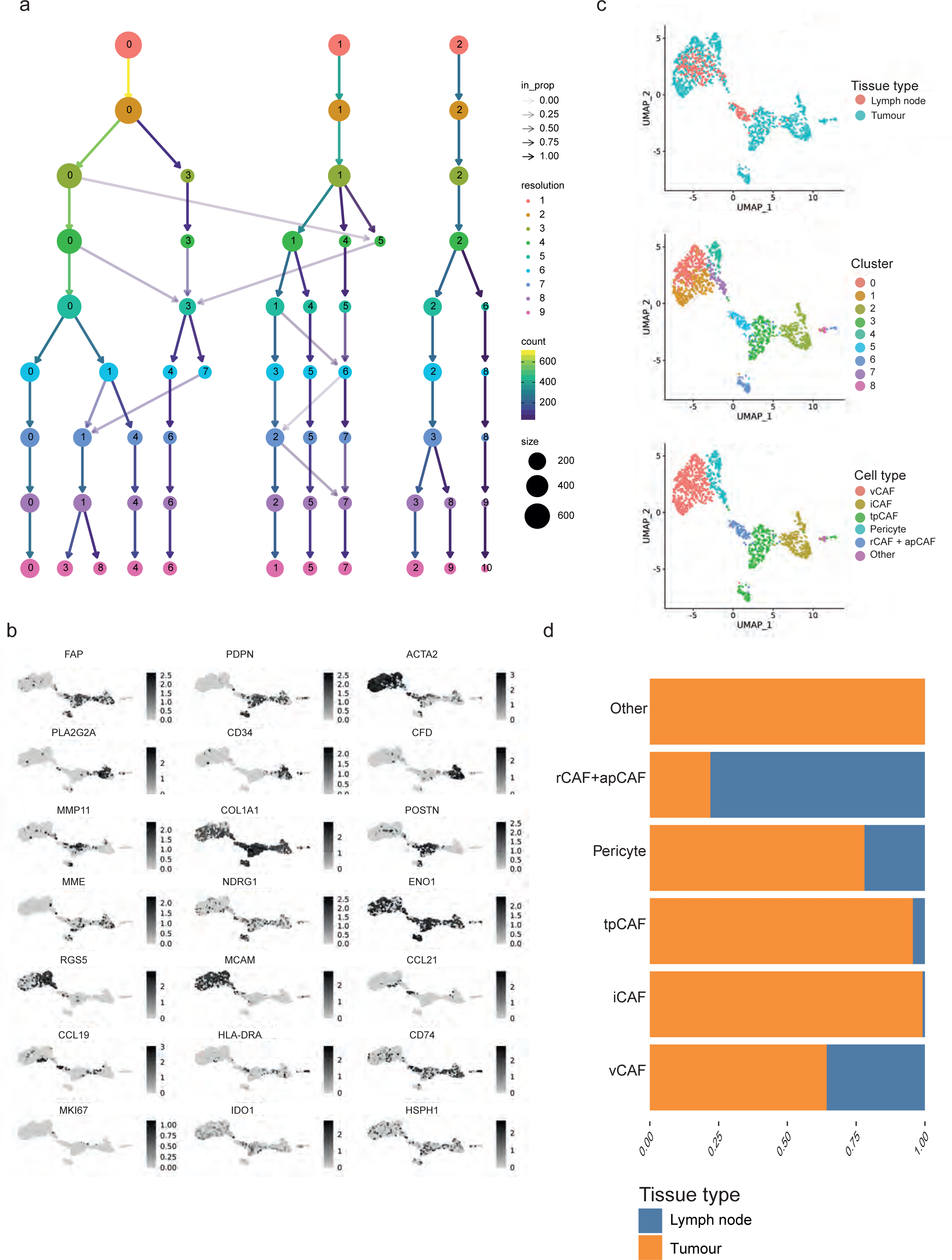
Analysis of scRNA-seq data from head and neck squamous cell carcinoma a) Clustree showing clustering hierarchy of HNSCC CAFs. Feature plot showing the marker expression of all marker genes for all cells in this dataset. b) UMAPs coloured by (top to bottom) tissue sites sequenced in the original study (metastatic lymph node, primary tumour tissue), clusters obtained at clustering resolution of 0.6, and the final cell type assignments of vCAFs, iCAFs, tpCAFs, pericytes, apCAFs + rCAFs, and other cells. c) Cell type distribution between tumour and metastatic lymph node tissue origin for all cell types defined in this dataset.

**Supplementary Figure 6.**
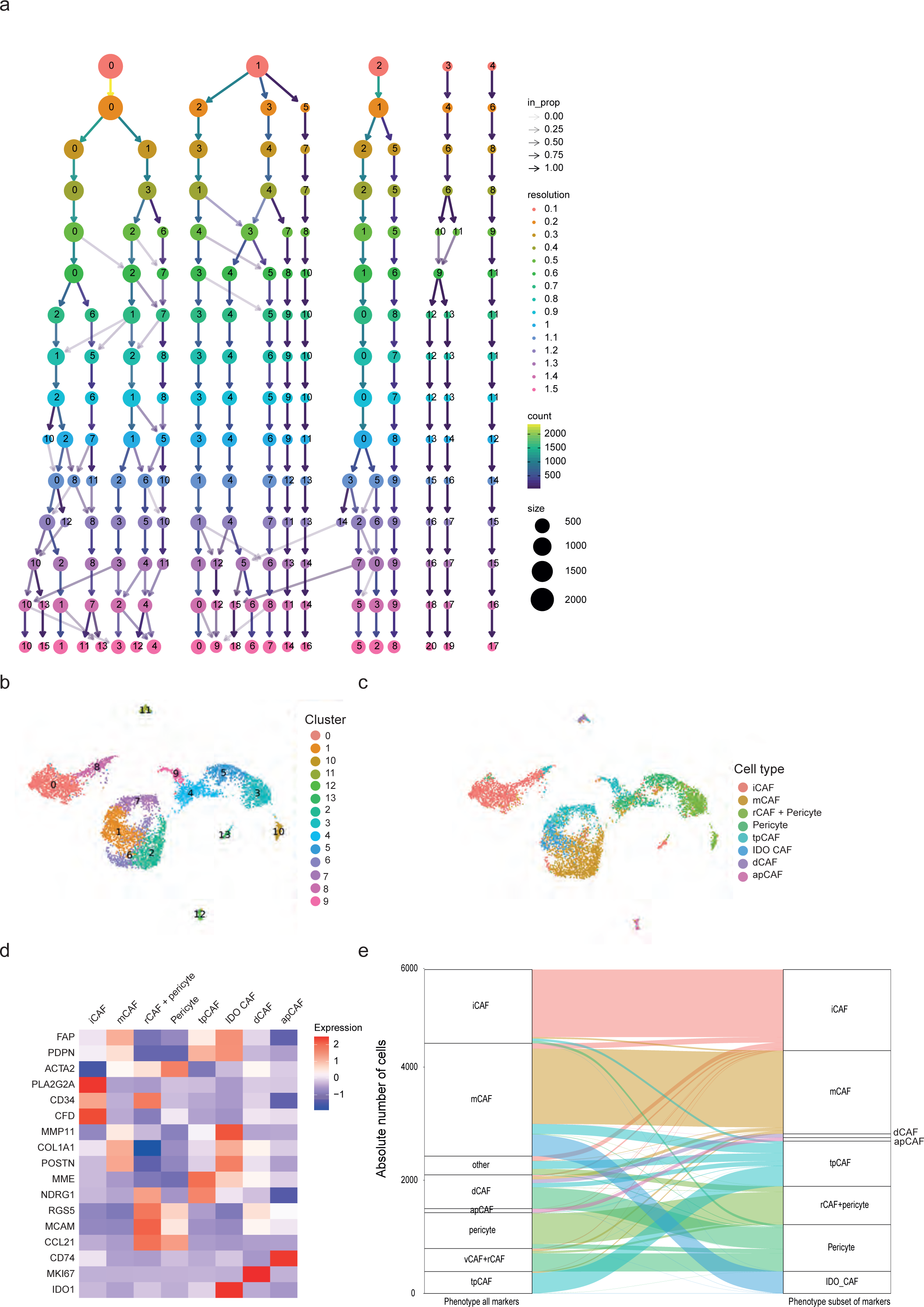
Analysis of the integrated scRNA-seq datasets a) Clustree showing the hierarchical clustering results of the integrated dataset. b) Clustering of the integrated dataset at 0.7 chosen for the final cell type assignment. c) UMAP coloured by CAF types recovered when clustering the integrated dataset using only the subset of defined marker genes. d) Heatmap showing the average expression level of defined marker genes for clusters defined after clustering the dataset using only the defined marker genes. e) Sankey plot showing the cell affiliations when clustering using (left) all genetic features of the integrated dataset and (right) only the subset of defined marker genes.

**Supplementary Figure 7.**
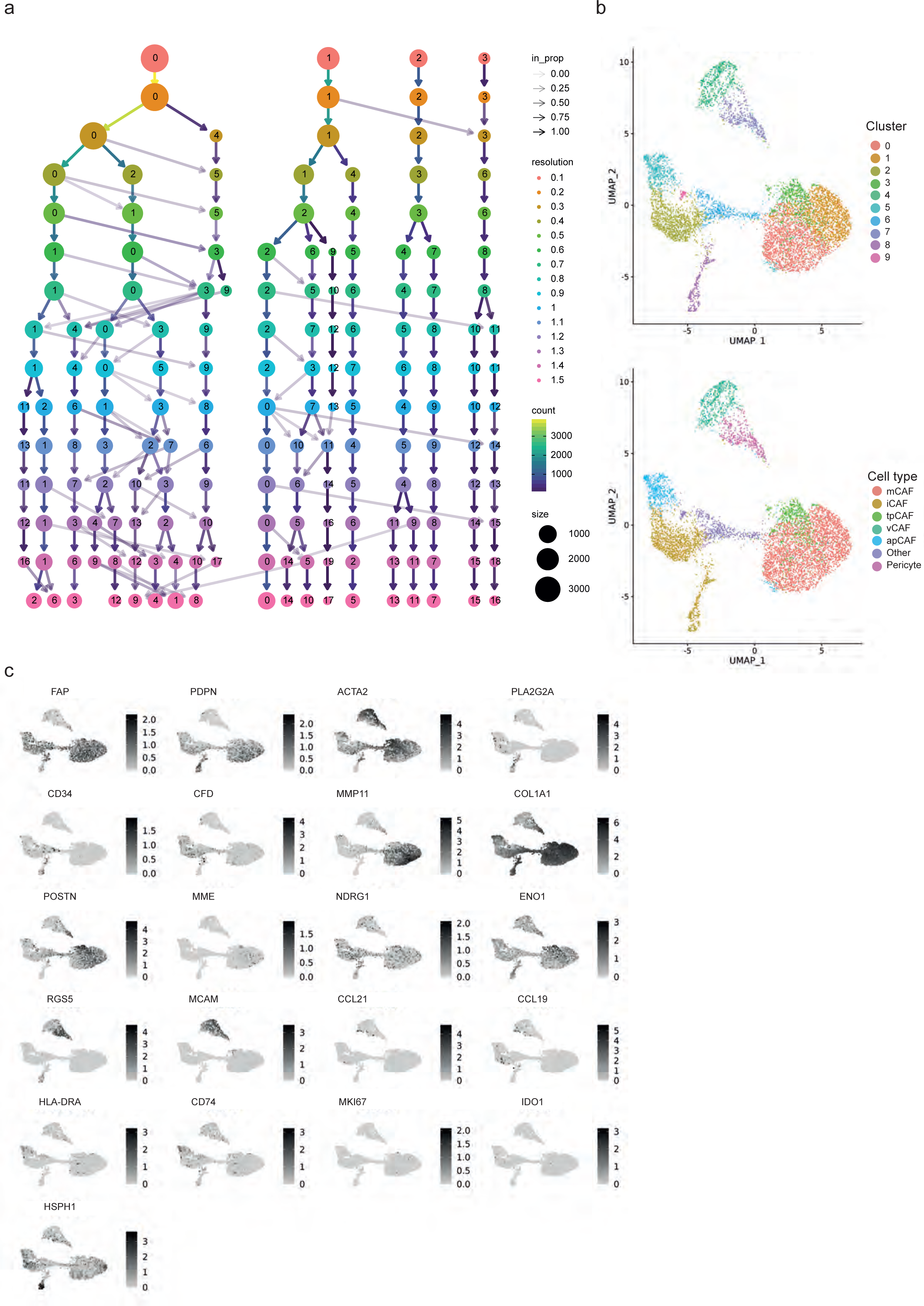
Individual analysis of a larger PDAC dataset a) Hierarchical clustering tree (using the clustree package) showing the clustering results of the big PDAC dataset. b) UMAPs coloured by the chosen clustering resolution of 0.6 and the resulting 10 different clusters as well as the final cluster assignment. c) Feature plot showing the expression of all previously identified marker genes on the UMAP.

**Supplementary Figure 8.**
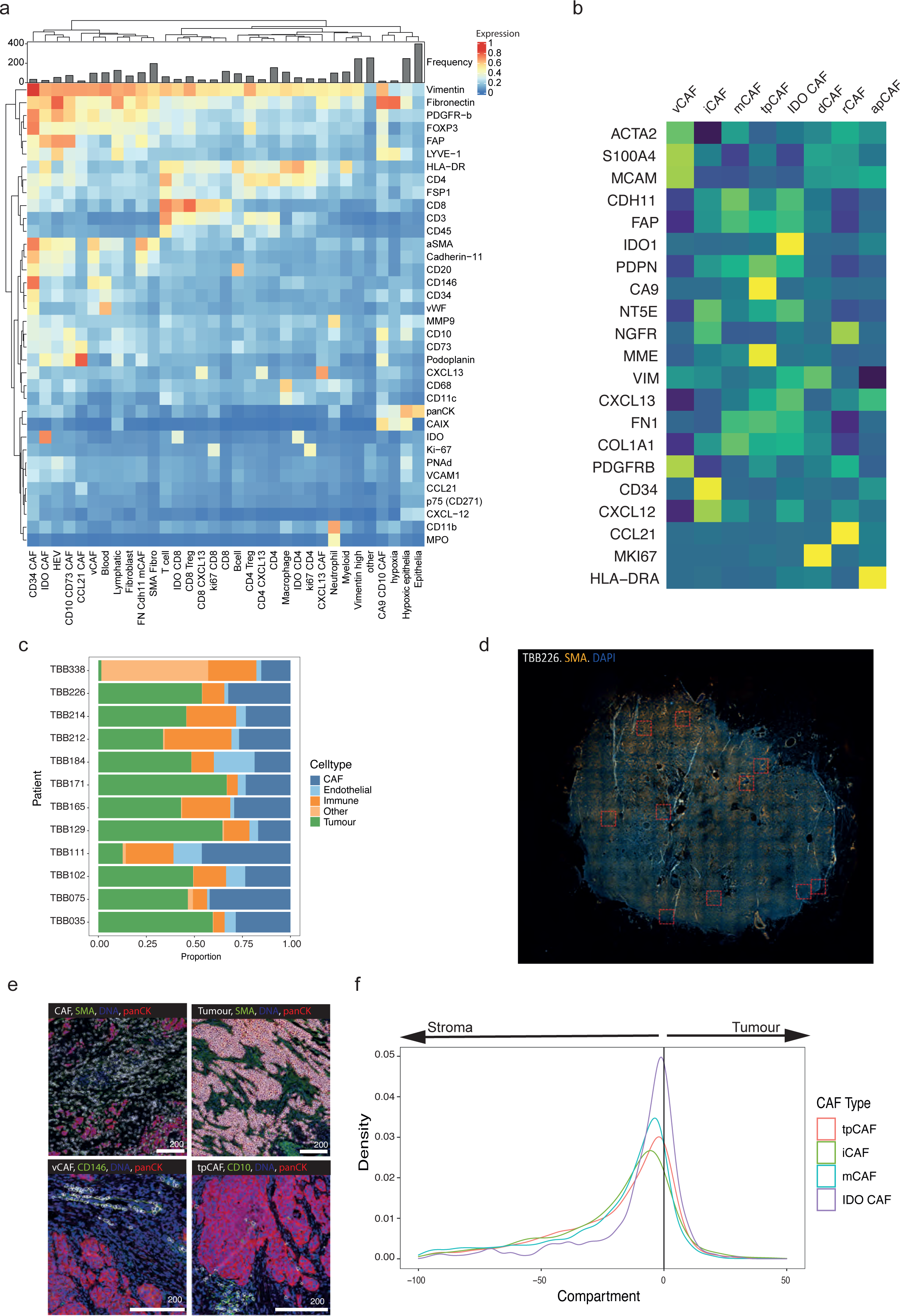
IMC analysis of matched breast cancer tissue samples a) Heatmap showing the average marker expression of all markers assessed in this study for all assigned cell types. Bar chart shows the number of cells per cluster. b) Average marker expression per CAF cluster in the breast cancer scRNA-seq data, when clustering based on the genes selected for IMC analysis. c) Bar chart showing the proportional distribution of CAFs, endothelial, immune, other cells, and tumour cells per patient as measured with IMC. d) Whole slide immunofluorescence image of sample TBB showing the region selection for IMC based on the stroma distribution. e) Images showing the expression of Iridium, SMA, panCK, CD146 and CD10 for a selection of images together with an overlay of cell masks showing the indicated cell types (all CAFs, tumour cells, vCAFs or tpCAFs) in the image. The images showing vCAFs and tpCAFs are zoomed-in areas to allow better visualisation of the CAF types in the tissue. Scale bars, 200 µm. f) Density plots showing the distance of CD34^+^ iCAFs, tpCAFs, mCAFs, and IDO CAFs to the tumour stroma border (stroma to the left, tumour on the right).

**Supplementary Figure 9.**
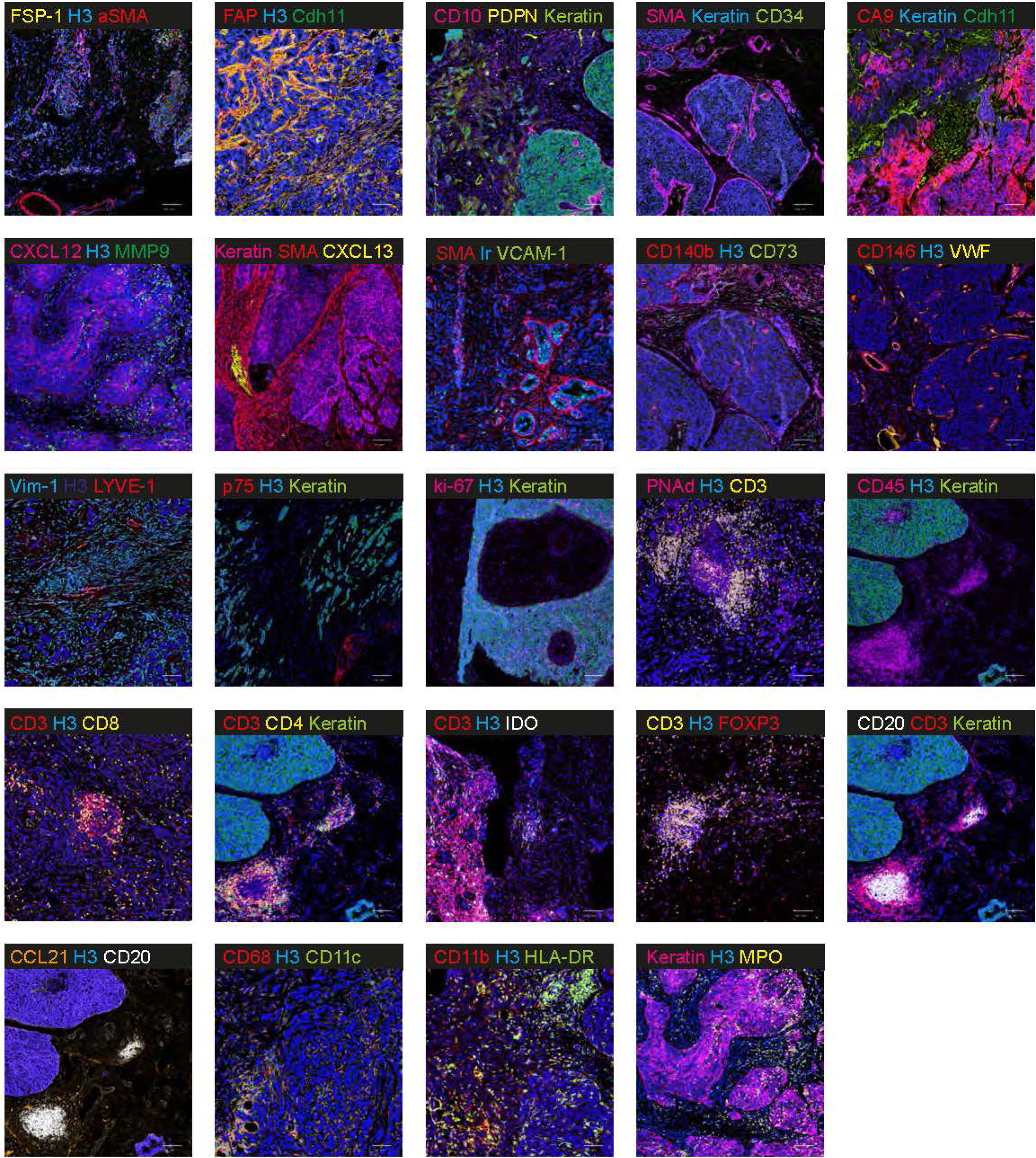
All measured IMC marker channels shown once All markers measured in IMC shown at least once as detected in tissues used in this study. Scale bars, 200 µm.

**Supplementary Figure 10.**
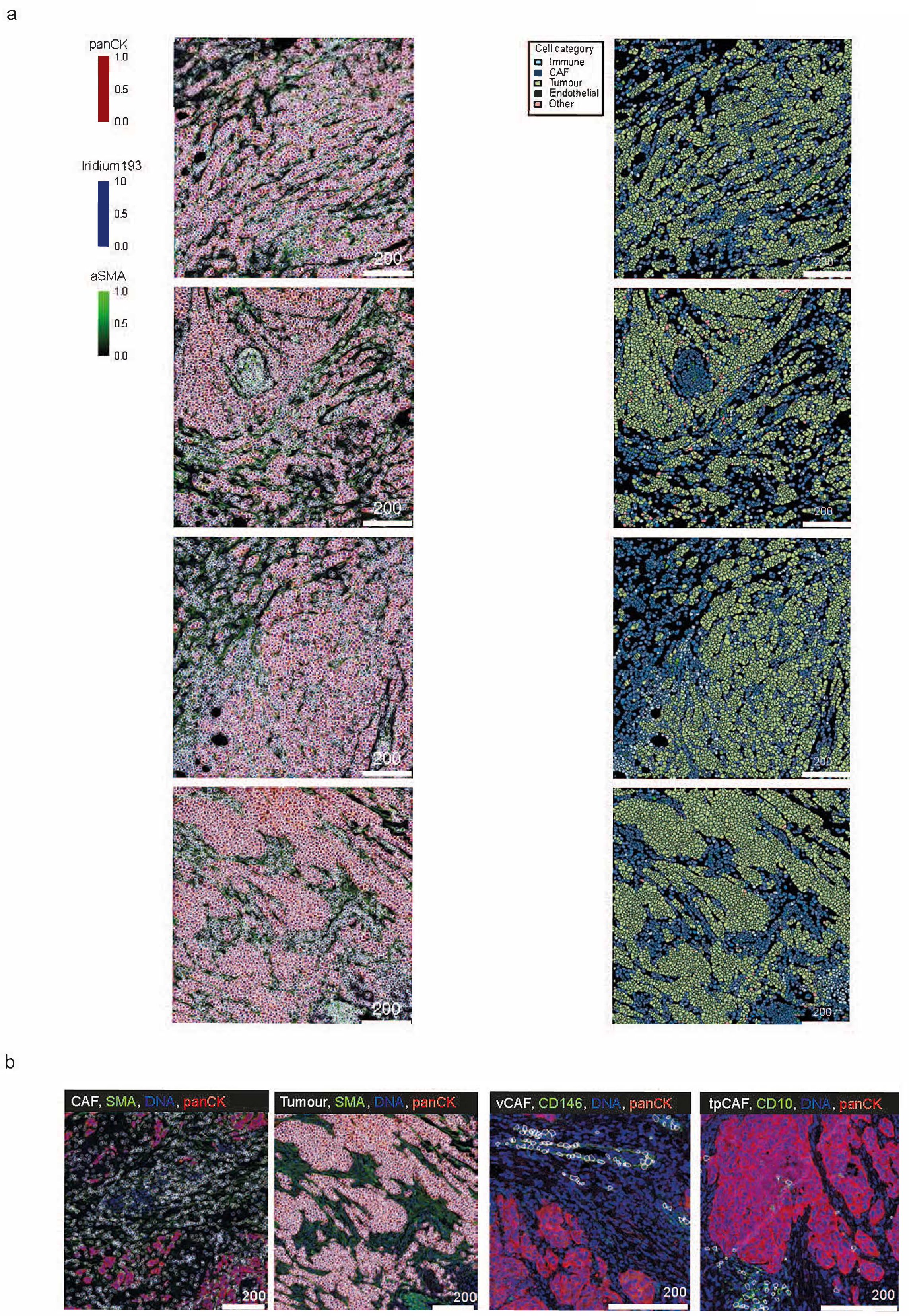
IMC single-cell segmentation mask overlays a) Raw pixel images (left) showing pan CK in red, SMA in green, DNA in blue with overlaying cell segmentation masks (cells in white). Pseudo-coloured segmentation masks (right) for all cell categories (tumour, CAF, endothelial, immune, other) defined in our IMC study. b) Raw pixel images showing expression of the indicated markers, with cell segmentation masks (white) also shown for CAFs, tumour cells, vCAFs and CD10+ CD73+ tpCAFs.

## Supplementary tables / Excel file titles and legends

Supplementary Table 1 – Clinical data and 10X run information Anonymised clinical data of all patients used in the breast cancer dataset.

Supplementary Table 2 – Differential gene expression analysis of breast cancer clusters as seen at resolution 0.4

Table showing results of differential gene expression analysis of all clusters as seen at resolution 0.4 before merging clusters to phenotypes

Supplementary Table 3 – Differential gene expression analysis comparing activated and non-activated CAFs

Table showing results of differential gene expression analysis of activated versus non-activated cells

Supplementary Table 4 – Differential gene expression analysis of all CAF and pericyte phenotypes

Table showing results of differential gene expression analysis of all defined CAF and pericyte phenotypes

Supplementary Table 5 – Differential gene expression analysis of all clusters as seen after dataset integration

Table showing results of differential gene expression analysis of all defined CAF and pericyte clusters in all datasets after dataset integration at the resolution of 0.8.

Supplementary Table 6 – Differential gene expression analysis of all phenotypes as seen after dataset integration

Table showing results of differential gene expression analysis of all defined CAF and pericyte phenotypes in all datasets after dataset integration

Supplementary Table 7 – Antibody panel and Research Resource Identifiers (RRID) validation

Table showing antibody panel, metal tags, antibody clone information and RRID validation information.

